# A systems biology approach to disentangle the direct and indirect effects of global transcription factors on gene expression in *Escherichia coli*

**DOI:** 10.1101/2022.04.18.488648

**Authors:** Mahesh S. Iyer, Ankita Pal, K. V. Venkatesh

## Abstract

Delineating the pleiotropic effects of global transcriptional factors (TFs) is critical in understanding the system-wide regulatory response in a particular environment. Currently, with the availability of genome-wide TF binding and gene expression data for *Escherichia coli*, several gene targets can be assigned to the global TFs albeit inconsistently. Here, using a systematic integrated approach with emphasis on metabolism, we characterize and quantify the direct effects as well as the growth-rate mediated indirect effects of global TFs using deletion mutants of FNR, ArcA, and IHF regulators under glucose fermentative condition. Apart from the previous known metabolite-TF interactions, we predicted with high confidence several novel interactions arising from the allosteric effects of the intracellular metabolites. Our combined experimental and compendium level computational analyses revealed the co-expressed genes regulated by these TFs as well as the coordination of the direct and indirect target gene expressions in the context of the economy of intracellular metabolites. Besides, we extended our analysis to combinatorial deletions of these TFs to determine their cross-talk effects as well as conserved patterns of regulatory interactions. Overall, this study leverages the fundamentals of TF-driven regulation which could serve as a better template for deciphering mechanisms underlying complex phenotypes.

## Introduction

Bacteria like *Escherichia coli* have developed complex regulatory strategies to sense and respond to environmental cues. The transcription factors (TFs), precisely the global ones, are the cornerstone in formulating an appropriate response (Babu et al., 2004; Grainger et al., 2005; Martínez-Antonio and Collado-Vides, 2003; Seshasayee et al., 2011). While the functions and the pleiotropic effects of the global TFs are well appreciated, the true direct targets and indirect effects on gene expression together with their underlying basis for a particular environment remain unclear. These genome-wide indirect effects are primarily mediated by alterations in growth rates or physiological state of the organism that encompass regulation by RNA polymerase associated sigma factors, ribosomes, intracellular metabolites as well as by other interacting global or local TFs (Berthoumieux et al., 2013; Gerosa et al., 2013; Klumpp et al., 2009; Li et al., 2019; Pal et al., 2022; Pan et al., 2021; Utrilla et al., 2016; Yu et al., 2021). For example, the growth rate was found to exert a significant effect on the gene expression patterns in a strain lacking the global regulator CRP (Pal et al., 2022). This growth rate-mediated effect on the gene expression was consistent in the event of adaptive laboratory evolution to facilitate the emergence of cells with better fitness (Pal et al., 2022; Utrilla et al., 2016). Thus, apart from the known functions of the direct target genes, it is fundamental to investigate the functions of these indirect or growth-rate mediated genes and their relevance in overall cellular objectives for a particular environment.

It is well known that around 51% of the gene expression changes are modulated by the global TFs (CRP, FNR, ArcA, IHF, HNS, Fis, and Lrp) while the remaining 49% are driven by their interaction with other pathway-specific local TFs (Martínez-Antonio, 2011; Martínez-Antonio and Collado-Vides, 2003). As gene expressions can be controlled by more than two regulators, it becomes crucial to determine the mode of interactions between the global TFs and other interacting or co-regulated TFs within the transcriptional regulatory network (TRN). Consequentially, understanding such interactions as to whether they stem from changes in gene expression of their interacting TFs or metabolite-driven changes in TF activities, remains fundamental. In a canonical sense, the transcript levels majorly dictate the abundance of enzymes followed by changes in metabolic flux and metabolite levels (Chubukov et al., 2014; Lempp et al., 2019; Link et al., 2013; Martin et al., 2012). The metabolite changes can often be attributed to the net effect of the metabolic flux as a result of the underlying proteome allocation (Iyer et al., 2021; Reznik et al., 2017; Scott et al., 2014; You et al., 2013). For instance, our previous studies have demonstrated the TF driven changes in proteome allocation that resulted in significant perturbations in the intracellular metabolite levels of the organism (Iyer et al., 2021; Pal et al., 2022). These intracellular metabolites not only constitute the biomass building blocks but also contribute as signalling molecules that confer metabolic plasticity to the organism (Chubukov et al., 2014; Erickson et al., 2017; Zampieri et al., 2019). *In vivo* studies concerning the allosteric effects of metabolites on TFs are now gaining importance as they further enhance our understanding of metabolism (Kochanowski et al., 2017; Lempp et al., 2019). However, identifying such metabolite-TF interactions during batch exponential growth based on steady-state physiological concentrations of the intracellular metabolites and activities of the TFs have been difficult. To determine the TF activities, one of the established approaches is the network component analysis (NCA) which works on a priori knowledge of the TF regulation on its cognate gene and gene expression values (Liao et al., 2003). A simple correlation analysis of the metabolite concentrations with the activities of TFs based on gene expression data will provide insights into the plausible metabolite-TF interactions relevant to glucose metabolism which can then be verified experimentally.

Apart from the metabolite-TF driven targets, other modes of regulation can also control the expression of the indirect or growth rate-mediated genes. Therefore, questions concerning the expression changes of specific sets of indirect genes, their cause and effect, still remain. Why do these gene expressions show perturbation in a specific pattern? Do the direct targets have a particular function that the organism is complementing using these indirect genes? Therefore, deconvoluting the complexity of gene expression in terms of co-expression as well as identifying the clusters of directly co-regulated genes becomes imperative (Tsai et al., 2018). In this regard, Weighted Gene Co-expression Network Analysis (WGCNA) represents one of the preferred algorithms. WGCNA involves the construction of modules of highly expressed genes based on similar gene expression profiles (Galán-Vásquez and Perez-Rueda, 2019; Langfelder and Horvath, 2008; W. Liu et al., 2018). These modules can be enriched using gene ontology analysis to identify the biological functions. However, the analysis of the modules in the context of maintaining the demand and supply equilibrium of intracellular metabolites remains largely unexplored. Moreover, given the availability of enormous gene expression datasets for *E. coli* K12 under various nutritional or stressful conditions (Meysman et al., 2014), it would be a valuable resource to derive the conserved co-expressed genes.

Here, we sought to investigate the complex layers of regulation driven by the global TFs using single and combinatorial knockouts of FNR, ArcA, and IHF and assessed the organisms’ response under glucose fermentative conditions. FNR and ArcA are best studied for their roles under anaerobiosis and energy metabolism (Federowicz et al., 2014; Grainger et al., 2007; Kang et al., 2005; Myers et al., 2013; Park et al., 2013). Similarly, IHF was been well characterized for its functions as a nucleoid protein as well as its role in central carbon metabolism (Kahramanoglou et al., 2011; Prieto et al., 2012). Our previous study elucidated the importance of these regulators in coordinating the expression of unnecessary or hedging genes, and necessary metabolic bottleneck genes together with intracellular metabolite adjustments required for optimal growth regime (Iyer et al., 2021). In this present study, we extended the examination of these global regulators with a focus on direct and indirect targets, interactions with other TFs, the co-regulated genes, and their preferences on metabolic genes towards formulating an appropriate response. Towards this, we utilized an integrated framework that disentangled the pleiotropic regulatory responses for a particular environment (Figure 1). Firstly, we determined the gene expression profiles and metabolite levels in the strains lacking these global regulators followed by extensive computational analyses. Next, we delineated the direct and indirect target gene sets and determined the regulatory activities and the corresponding metabolite-TF interactions which altogether gave us insights into the regulatory interactions. Moreover, we have also elucidated the coordination or co-expression between the direct and indirectly coregulated genes by employing WGCNA on *E. coli* K12 compendium gene expression data. Furthermore, we elucidated the crosstalk between these global regulators using combinatorial deletions of these TFs to decipher whether they exhibit additive or non-additive effects on the genes that they regulate. Overall, our study can serve as a template for understanding systems-level regulatory events that are even applicable for deciphering the mechanistic strategies developed during pathogenesis, gut microbiome interactions, and cancer metabolism.

**Figure 1.**
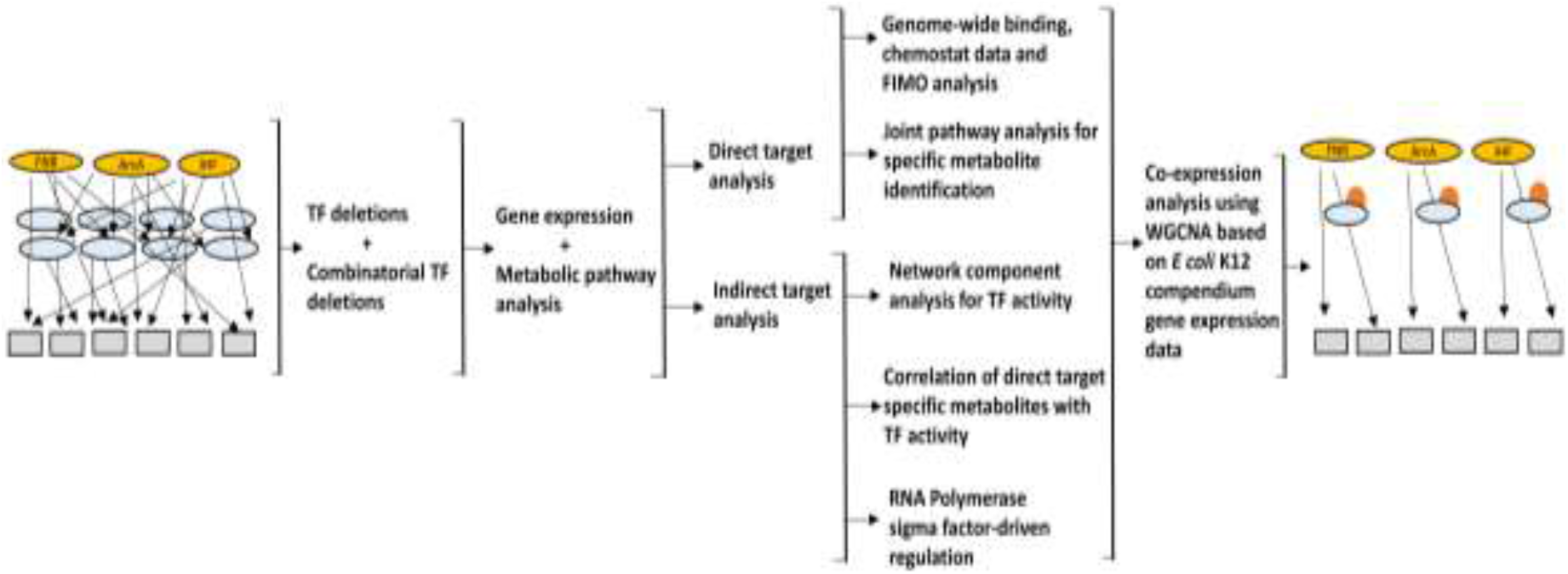
An overview of the integrated workflow to dissect the complex regulation on gene expression existing within the TRN. In this study, these analyses were carried out for the global regulators FNR, ArcA and IHF by employing single or combinatorial deletions in anaerobic glucose fermentative conditions. In this figure, the yellow filled circles represent global TFs, the blue filled circles represent other TFs (global or local),the orange filled circles represent metabolites, the grey boxes represent the direct and indirect metabolic gene targets and the arrows represent regulations (positive or negative).

## Results

### Crosstalk and nature of interaction within FNR, ArcA, and IHF regulators

We constructed three double mutants FA (Δ*fnr*Δ*arcA*), FI (Δ*fnr*Δ*ihf*), and AI (Δ*arcA*Δ*ihf*), by deleting *arcA, fnr*, and *ihf* (both *ihfA* and *ihfB*) genes in combination, using homologous recombination (Datsenko and Wanner, 2000). We monitored the gene expression profiles for the double mutants compared to WT in the mid-exponential phase with glucose minimal media. The gene expression data observed in FA mutant was found to be consistent with a previous study (Covert et al., 2004) (Supplementary File S1). However, the gene expression of the mutants FI and AI have not been characterized so far. The gene expression data for the single mutants were obtained from our previous study (Iyer et al., 2021). We investigated the mode of interaction between each of the regulators in the double mutant, specifically, whether the expression patterns could be explained by either of the deleted TFs or by combined pleiotropic effects of both the regulators, defined hereafter as ‘crosstalk’. Preliminary analysis was performed for both the double and single mutants using the total number of differentially expressed genes (DEGs, P < 0.05, Benjamini-Hochberg (BH) adjusted), log2 fold change >= |1|). We observed ~250, ~510, and ~1250 DEGs in FA, FI, and AI mutant compared to WT, respectively. In FA and FI mutants the number of DEGs were however lower than the DEGs in their single mutants combined (Figure 2A). This was in contrast to AI wherein the number of DEGs was greater than the sum of the DEGs in its single mutants. This indicated a greater extent of crosstalk existing between ArcA and IHF regulators than other combinations involving FNR.

**Figure 2.**
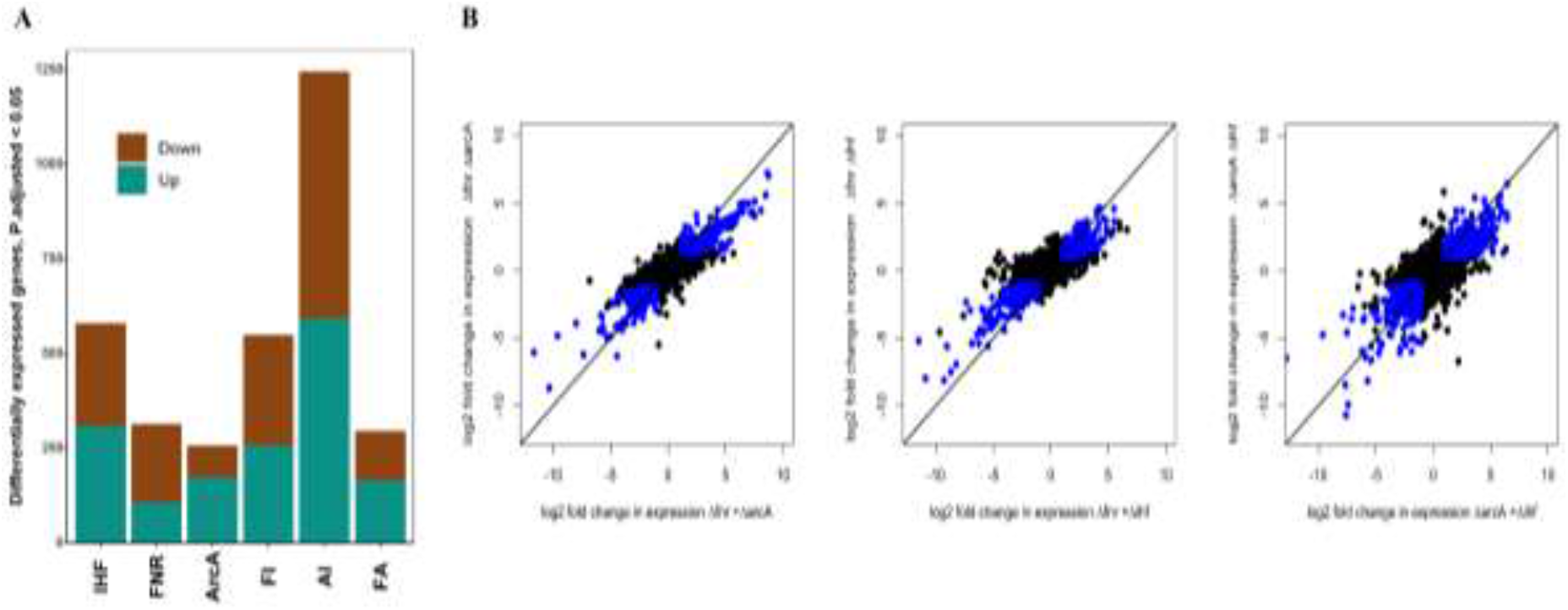
Transcriptome comparison of the regulator mutants compared to WT. **A)** Bar plot represents the number of upregulated and downregulated DEGs in each of the single and double mutants compared to WT. **B)** Scatterplot depicting the mode of interaction within FNR, ArcA and IHF regulators using log2 fold changes in the double mutant compared to the sum of log2 fold changes in the single mutants, respectively. The blue dots represent DEGs in double mutants (≥ 2 fold) and genes having fold change ≥ 2 fold by summing the log2 fold changes in each of its single mutant.

To understand the exact nature of these crosstalk as well as to gain quantitative insights, we first assessed whether the DEGs in the single mutants reproduced their gene expression patterns in the double mutants. We found that the genes whose expression were significantly downregulated or upregulated (DEGs, FDR < 0.05, log2 fold change ≥ |1|) in the single mutants, were also downregulated or upregulated in the double mutant (log2 fold change < 0 or > 0) (Supplementary Information, Figure S1 & S2). Next, we plotted the genes which were differentially expressed in the double mutant but showed more than 2-fold change in gene expression in both single knockouts added together (Figure 2B). The results provided direct evidence of widespread additive effects between each of the regulators, as either of the single mutants could explain the observed changes in the double mutants. We quantified the effects of each of the regulator deletions in the double mutants and performed a two-proportion Z-test with Yate’s continuity correction for assessing significance (P < 0.01, Supplementary File S1). In the DEGs of the double mutants, upregulated or downregulated gene expression changes could be attributed to specific regulator effects rather than a collective effect. Besides, the genes which could not be assigned (~20-38%) to either regulator deletion effects were significant in the case of AI followed by FA and FI mutants, depicting the extent of crosstalk. Among DEGs of the double mutants, we observed only ~11-20% overlapping genes of both the single mutants with more than 2-fold change in gene expression. Overall, these data indicated that in the double mutants, each of the regulators showed independent but additive crosstalk effects due to their deletion.

### Regulatory effects on key metabolic pathways

In our previous work, we demonstrated the systems-wide effect of the transcriptional regulators FNR, ArcA, and IHF on key metabolic pathways (Iyer et al., 2021). To decipher whether those metabolic pathways were perturbed in the double mutants as well, we carried out a KEGG pathway enrichment analysis (Proteomaps (Liebermeister et al., 2014) based gene classification) on DEGs obtained in double mutants with statistical significance analysis. Of the total upregulated and downregulated DEGs in FA mutant, 74% and 69% of genes showed enrichment for metabolic pathways, respectively (Supplementary File S1). In these KEGG pathway enriched (KPE) upregulated DEGs, we observed significant enrichment of amino acid metabolism related to arginine degradation that generates internal ammonia, costly TCA cycle genes unnecessary under anaerobic fermentation, and unneeded lipid and steroid degradation genes (Figure 3A). Additionally, in KPE downregulated DEGs we obtained significant enrichment of amino acid metabolism and transport-related genes (Figure 3B). The amino acid metabolism genes belonged to arginine and asparagine biosynthesis whereas the transporters belonged to either alternate or secondary carbons. In the case of FI mutant, 57% and 58% of downregulated and upregulated genes showed enrichment towards metabolic pathways, respectively (Supplementary File S1). The KEGG pathway downregulated DEGs (Supplementary Information, Figure S3) showed enrichment for amino acid metabolism related to branched-chain amino acid biosynthesis, glycolysis, chaperone folding and catalyst-related pathways. Alternatively, only transport encoding genes were upregulated in KEGG pathway enriched DEGs. On the other hand, 53% and 66% of down-regulated and up-regulated genes showed enrichment towards metabolic pathways in AI mutant, respectively (Supplementary File S1). We obtained enrichment for amino acid metabolism related to branched-chain amino acid biosynthesis and bacterial motility proteins in downregulated KEGG pathway enriched DEGs in AI mutant compared to WT (Supplementary Information, Figure S4). Amino acid metabolism related to putrescine and arginine degradation, aerobic oxidative phosphorylation genes and TCA cycle genes, transporters, and several transcription factors were upregulated in KEGG pathway enriched DEGs in AI mutant compared to WT. These data indicating significant perturbations of metabolic pathways in the double mutants were consistent with the observations in single mutants. Indeed, we were able to demonstrate the shared patterns of regulation along the metabolic pathways in the double mutants to either of its single mutants. For instance, the metabolic pathway analysis in FA mutant showed dominant effects of ArcA whereas both FI and AI were strongly inclined towards IHF. Besides, the pattern of enrichment was also consistent with previous observations in the case of single mutants wherein the necessary metabolic pathways of amino acid and nucleotide metabolism were downregulated and unnecessary metabolic pathways such as the aerobic TCA cycle, oxidative phosphorylation, and alternate carbon transporters were upregulated.

**Figure 3.**
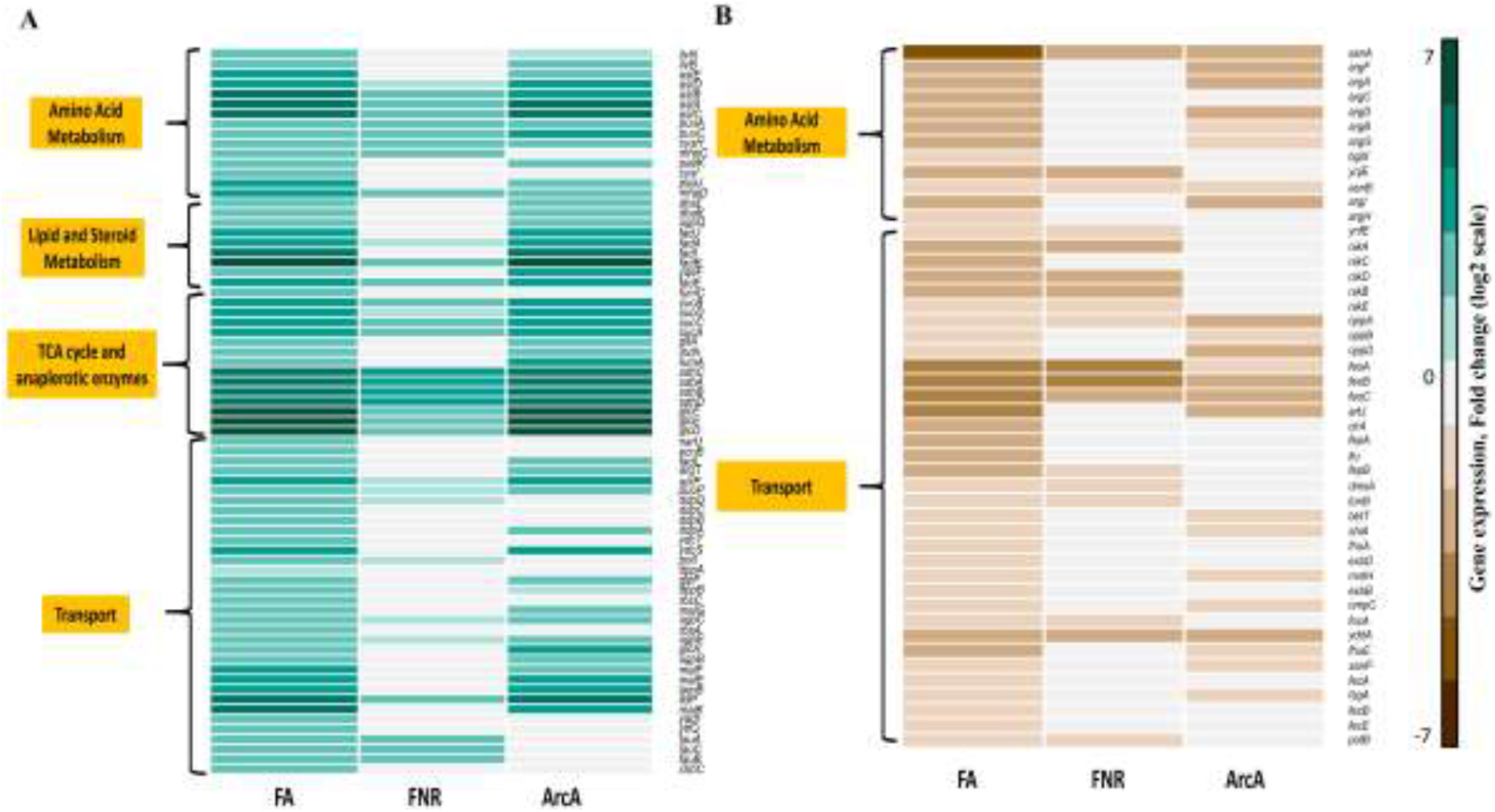
Heatmap depicting the metabolic pathway analysis of regulator mutants compared to WT. The figure shows the comparison of **A)** upregulated and **B)** downregulated DEGs of FA mutant compared to WT. Only the significantly enriched KEGG pathways are shown. For comparison across the strains, only the DEGs found in FA, Δ*fnr* and Δ*arcA* mutant compared to WT were considered. See Figure S3 and S4 for metabolic pathway analysis of DEGs in FI and AI respectively.

### Loss of TFs results in changes in growth rates and metabolite levels

We monitored the physiological effects namely glucose uptake, growth rate, and secretion profiles of mixed acid fermentation products as a result of the deletion of TFs. We observed detrimental effects on growth rate in all the double mutants relative to WT (Supplementary Information, Table S1). The mutants FA, FI, and AI showed a reduction in growth rate by 40%, 30%, and 37% respectively. This reduction in the growth rates was consistent with the severe perturbation seen in the metabolic pathways. The glucose uptake reduction in FA and FI mutants strongly correlated with their respective growth rate changes. Interestingly, the reduction in glucose uptake in AI mutant was only 13% compared to WT, but not as severe as its reduction in growth rate. A severe reduction in biomass yield was also observed in the case of AI mutant (23%) indicating reduced carbon partitioning towards biomass (Supplementary Information, Table S1). The secretion profiles of exo-metabolites namely ethanol, acetate, formate, lactate and succinate revealed distinct changes in their yields (Supplementary Information, Table S1). Towards a detailed understanding of metabolism, we monitored the intracellular metabolite concentrations of the glucose metabolism in all the mutants using ^13^C labelled targeted metabolomics (Iyer et al., 2021; Pal et al., 2022). Among the measured metabolites, the intracellular concentration of glycolytic and TCA cycle intermediates and proteinogenic amino acids were significantly different compared to the WT (Student’s t-test, FDR < 0.05). In FI and AI, we observed increased accumulation of the metabolic precursors and amino acids consistent with the accumulations seen in the single mutants as reported previously (Iyer et al., 2021) (Supplementary Information, Figure S5). These accumulations reflected the inability of the mutants to utilize the metabolites towards biomass synthesis, in agreement with their reduced growth rates. However, we observed lowered pool sizes of these metabolites in the FA mutant compared to WT. This might indicate an adverse carbon limitation (Tweeddale et al., 1998) as well as the reduced availability of active ribosome pools (Dai et al., 2018, 2016; Li et al., 2018; Metzl-Raz et al., 2017; Scott et al., 2014) prevalent in slow-growth conditions, which needs to be investigated in the future.

### Dissecting direct and indirect targets of the TF from their pleiotropic effects on metabolism

Global TFs have direct as well as indirect pleiotropic effects on gene expression. The direct effects can be attributed to the condition-specific binding of the TFs to the upstream sequences of the target gene. On the other hand, the indirect effects are primarily the growth rate-mediated effects (Li et al., 2019; Pal et al., 2022; Utrilla et al., 2016) arising from the deletion of the TFs, in this case, the absence of either FNR, ArcA and IHF and their combinatorial deletions, referred hereafter as dTF/s. Therefore, we sought to disentangle the growth rate-mediated effects from the direct effects of the dTFs. Here, we employed a multi-pronged approach using the growth-rate dependent genes obtained from chemostat, binding studies available to enrich for the binding region of the regulators, and comparison with available gene expression data to account for all the genes modulated purely by the global dTF. First, we performed glucose-limited chemostat cultivations of the WT strain under aerobic respiratory and anaerobic fermentative conditions at a fixed dilution rate of 0.21 h^-1^ to identify the genes whose expression changes were growth-rate independent. A comparison of DEGs for WT in anaerobic fermentation with WT in aerobic conditions performed under the chemostat conditions represent the genes that are specific to regulation or genes that are not altered due to slow growth effects in anaerobic fermentation. The genes that were not differentially expressed in the glucose-limited chemostats were attributed to lowered growth rates. It has been shown previously that the gene expression changes in response to growth rate effects are consistent, despite the glucose-limited or glucose-excess conditions (Li et al., 2019; Pal et al., 2022; Utrilla et al., 2016). Thus, the DEGs identified under the chemostat conditions were then utilized to characterize the indirect targets as growth-rate mediated effects and the direct targets as regulation-specific effects by the dTF/s in batch-exponential growth conditions. The DEGs enriched for direct targets of the dTFs in the double mutants were further validated by determining the binding motifs using FIMO analysis (Charles E Grant et al., 2011) together with the available *in vivo* ChIP binding or other *in vitro* binding data (Federowicz et al., 2014; Grainger et al., 2007; Keseler et al., 2017; Myers et al., 2013; Park et al., 2013; Prieto et al., 2012; Santos-Zavaleta et al., 2019). Further, a few regulation-specific genes (15-20%) in double mutants and (20-32%) in case of single mutants, found in our analysis that were not characterized as direct targets, could potentially represent uncharacterized novel targets (Choudhary et al., 2020) of the dTF/s.

We delineated the direct regulation-specific or indirect growth-rate mediated effects on metabolism by taking into consideration the KPE DEGs. Overall, we found that 55-60% of upregulated and downregulated KPE DEGs to be regulation-specific in FA compared to 30-33% in FI and AI mutants (Figure 4A). The slow growth mediated changes in gene expression were dominant in FI and AI mutant. In the case of single mutants, the proportion of indirect genes were prominent in Δ*ihf* followed by Δ*fnr* and least in Δ*arcA*. The significantly enriched pathways within the upregulated KPE DEGs in the double (except FI) as well as the single mutants (except Δ*ihf*), showed a greater extent of regulation-specific effects. However, the data did not ascertain whether the reduction in growth rate was proportional to the extent of growth rate-mediated or indirect targets. On the other hand, the significantly enriched pathways within the downregulated KPE DEGs of the strains (except FA) aligned well with the reduced growth rate (Figure 4B-D, Supplementary Information, Figure S6A-C).

**Figure 4.**
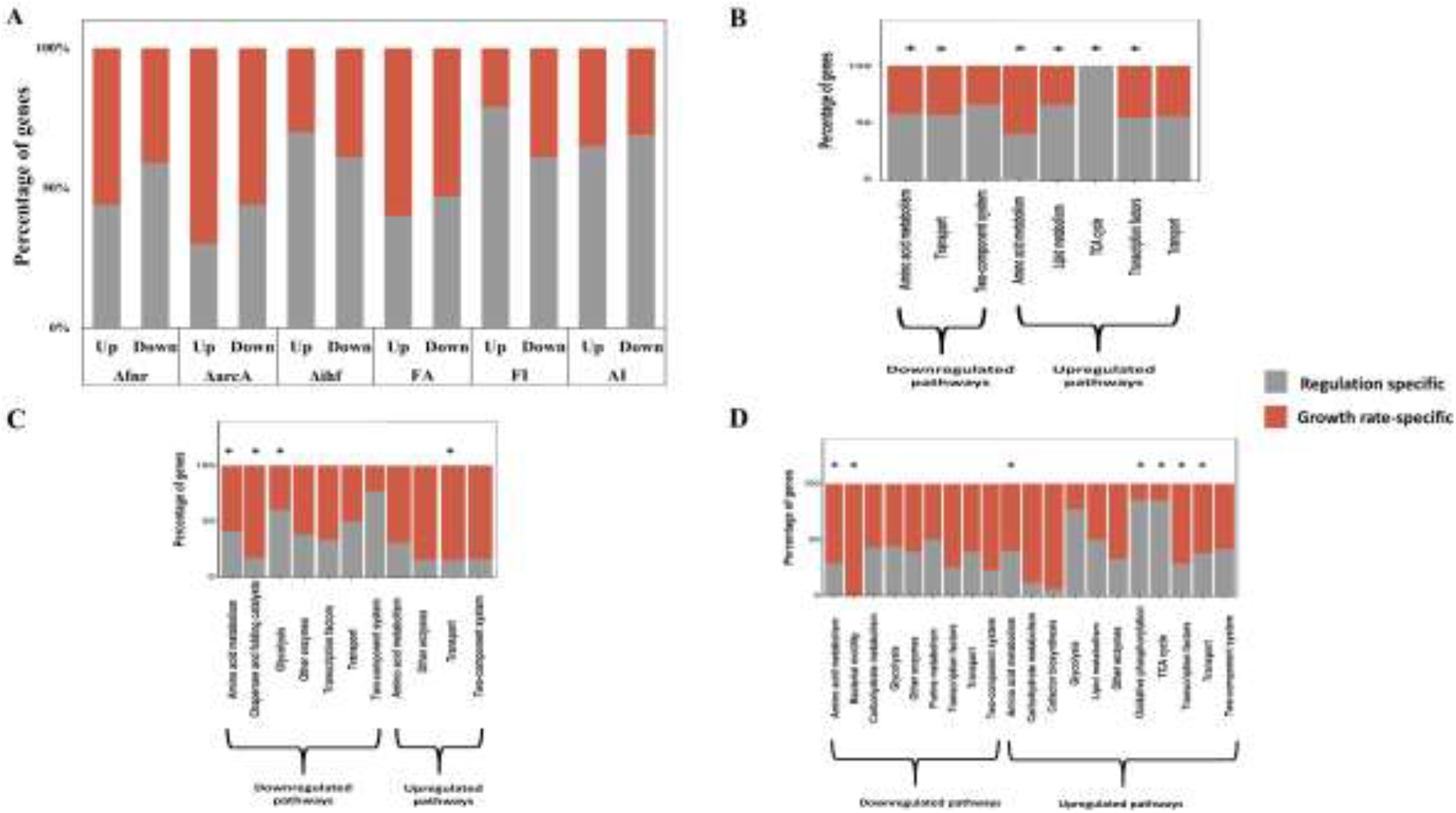
Direct and indirect effects on gene expression. Stacked plot depicting the percentage of direct targets and growth-rate mediated indirect targets **A)** in KPE DEGs, as a result of deletion of the global TFs **B)** FA, **C)** FI and **D)** AI compared to WT. Asterisk represent the significantly enriched metabolic pathways. Only the metabolic pathways which had at least 9 genes were retained for the analysis. See Supplementary File S1 for the complete list. See Figure S6 for the single TF-deleted mutants.

### Exploring the regulatory interactions of dTF/s with the iTFs on the indirect gene target expressions

Next, we analysed the indirect gene targets for any TF enrichment other than the dTF, assuming the probability of regulating these genes by the dTF is low or they might represent silent targets (Choudhary et al., 2020; Park et al., 2013) in this particular condition. These TFs are other interacting TFs (referred hereafter to as iTFs) whose gene expression or activity are modulated by the dTF/s. The enrichment of genes for the presence of binding from iTFs was done using EcoCyc or RegulonDB (Keseler et al., 2017; Santos-Zavaleta et al., 2019) and further assessed for statistical significance (Fischer exact test, FDR < 0.05).

We postulate that enrichment of the direct gene targets majorly responded to the changes as a result of the dTF/s. We observed in the upregulated and downregulated indirect targets the enrichment of global as well as local iTFs (Supplementary File S1). The activity of the iTFs was determined based on the manual interpretation of the changes in its target gene expression. For instance, we defined the activity of an iTF to be decreased when there was de-repression of its gene targets. We found that the iTFs enriched in the upregulated genes were unique from the downregulated genes for each of the strains. Further, we observed that PuuR iTF activity was perturbed in all the strains (except FI) followed by Crp (in 4 of 6 strains), Fur (3 of 6 strains), and CysB (3 of 6 strains). PuuR is the regulator that represses the expression of putrescine degradation required for the generation of ammonia and response to various stresses. In most of the cases, changes in activity of CRP and PuuR iTFs were either not correspondent or antagonistically correspondent with the changes in gene expression of *crp* or *puuR* (except in FI where reduction in CRP activity was concomitant with a reduction in *crp* gene expression). Only MhpR in AI, Nac in FI, and ArcA in Δ*ihf* showed an increase in their gene expression concomitant with their increased activities. Overall, assessment of the changes in gene expressions of the iTFs itself revealed that the activity of the iTFs was perturbed due to reasons not involving its gene expression. The other possible mechanisms involve changes at the post-translational level or the protein level. As the transcript abundances majorly dictate the protein concentrations (Schmidt et al., 2016; Yu et al., 2020), we continued our investigation focusing on the dTF-iTF interaction at the post-translational level.

The metabolic precursors and amino acids, apart from being building blocks for protein biomass, can also act as signalling molecules that can allosterically modulate the activities of the TFs to bring about changes in the gene expression of the organism (Chubukov et al., 2014; Lempp et al., 2019; Link et al., 2013; Reznik et al., 2017; Zampieri et al., 2019). The concept to define dTF-iTF interaction was that metabolites perturbed due to changes in direct targets by dTF/s were allosterically regulating the iTFs which then modulated the indirect or growth rate mediated gene targets. These metabolites that responded to changes in direct targets were mainly glutamate, aspartate, PEP, αKG, GABA, citrate, glycine and to a lesser extent glutamine, arginine and lysine. The other metabolites which were significantly (FDR < 0.05) perturbed in each of the strains could be ascribable to the concomitant changes in its precursor metabolites. For instance, changes in methionine levels were concomitant with the changes in its precursor aspartate (Supplementary Information, Figure S5).

To address the putative allosteric metabolite-iTF interactions, we carried out a linear correlation (Pearson correlation coefficient, r > 0.75, P < 0.05) between the KPE direct target specific metabolite concentrations and the iTFs activities. As metabolite-iTF interactions by linear correlation are prone to high false positives and to increase the confidence of such putative metabolite-iTF interactions, two stringent criteria were applied. Firstly, metabolites that were perturbed due to the direct targets of dTFs were only considered. To obtain the list of metabolites, a joint pathway analysis was performed using the significantly perturbed metabolites with the combined upregulated and downregulated KPE direct targets for each strain separately and metabolite-gene pair which satisfied the minimum threshold (see Methods) were retained for that particular strain. Secondly, the metabolite-iTF pair was reanalyzed using the pathway information available in EcoCyc to identify whether the metabolite (Pearson correlation coefficient, r > 0.9, P < 0.05) is involved in any way with the KPE indirect gene targets of that particular iTF. This works similar to the distance criteria (Lempp et al., 2019) approach wherein the iTF regulating the gene (encodes enzyme) and metabolite are part of the same metabolic subsystem. A further addition to the analysis involved the comparison of the metabolite-iTF interactions with published studies reporting similar putative metabolite-TF interactions (Lempp et al., 2019).

To quantify the iTF activities, we recalled the network component analysis (NCA) algorithm (Liao et al., 2003) that derives the regulatory activity from the experimentally measured gene expression profiles. NCA has been previously used to study large scale gene regulatory networks (GRNs) and successfully reproduces majority of the activity profiles of the TFs depending on the mode of interactions with its target genes (Lempp et al., 2019). For each of the strains, NCA was performed specifically on the KPE growth-rate mediated or indirect genes. Only those iTF activities that NCA was able to reproduce identical to the manual interpretation, were considered for metabolite-iTF interactions analyses. Further, if the iTF activities were deemed unclear from the manual interpretation, in the sense, those that cannot be defined as positive or negative (<70% targets, dual role) regulators were excluded from NCA analysis. With NCA, we were able to reproduce 80% of the iTFs activities for each of the strains including the known interactions as well as novel putative metabolite-iTF interactions (Supplementary Information, Figure S7A-F). For instance, we were able to reproduce the known interactions of PuuR with putrescine (inferred from GABA (Iyer et al., 2021)) as well as Crp with cAMP (inferred from PEP (McCloskey et al., 2018; Pal et al., 2022)). The metabolite cAMP was not detectable in our LC-MS run probably due to the limit of detection and anaerobic fermentation conditions with high glucose uptake rate (was detectable in aerobic condition). As cAMP and PEP levels are interdependent (Deutscher et al., 2006; Postma et al., 1993), we have assumed the association of Crp with PEP similar to its association with cAMP. ln FA mutant, we observed increased activity of Crp, NtrC and Fur and decreased activity of PuuR along with their allosteric effectors such as PEP, αKG, citrate, GABA, glycine and arginine (Supplementary File S1). This was consistent with increased activity of Crp with its interacting metabolites PEP, αKG and citrate in Δ*arcA* and NtrC in Δ*fnr* as well as decreased activity of PuuR with its allosteric metabolites GABA, PEP and αKG in both the single mutants. In FI mutant, we observed increased activity of Fur, CysB and Nac and decreased activity of Lrp. The activity changes of CysB and Lrp along with their interactions with metabolites GABA, arginine, glutamate, αKG and PEP were consistent with the activity profile observed in the Δ*ihf* mutant. In AI, we observed a large number of regulators with changes in their activity profiles, concomitant with the widespread changes in the gene expression profiles which reinforces the greater crosstalk effects between ArcA and IHF. The decreased activity of Lrp, PuuR, TrpR and UlaR and the increased activity of CysB along with their interactions with the metabolites’ glutamine, glutamate, aspartate, PEP, arginine and GABA in the AI mutant was consistent with the activity profiles of Lrp, CysB, TrpR and UlaR in Δ*ihf* and PuuR in Δ*arcA* mutant.

We examined across the strains the metabolite-iTF interactions as a function of growth rate (Figure 5A). This analysis not only revealed the shared patterns of interactions seen in the double and single mutants but also the crosstalk effects. For instance, occurrence of metabolite-ArcA were more prominent towards higher growth rates as opposed metabolite-Fur seen towards lower growth rates. AI showed the highest number of metabolite-TF interactions compared to its single mutants which reiterated the crosstalk effects in contrast to the interactions seen in FA and FI mutants. Based on all the metabolite-iTF interactions across the strains, we wanted to determine the conserved metabolite-iTF pairs. In total, we observed 119 connections between 10 metabolites and 16 iTFs (Figure 5B). It should be noted that we observed enrichment of ArcA in both Δ*ihf* and Δ*fnr* mutants as a result of which ArcA was considered as an iTF in the metabolite-iTF interaction analysis. We found that 12 metabolite-iTF connections (glutamate-ArcA, glutamate/GABA/arginine-PuuR and αKG/glutamine-Fur) which showed positive as well as negative associations accounted for only 10%, whereas remaining 90% of the metabolite-iTF connections were unique. We observed that glutamate, aspartate, PEP and GABA metabolites had the greatest number of allosteric connections with iTFs. They were followed by metabolites arginine, α-ketoglutarate, and glutamine with medium connectivity and glycine, lysine, and citrate with low connectivity with their respective iTFs. Next, based on the stringent criteria described above, we ended up with a total of 29 connections with 9 iTFs and 8 metabolites (Figure 5C). Among these, ~45% of the interactions/associations were reported in a previous study (Lempp et al., 2019). Overall, we were able to determine with high confidence novel metabolite-TF interactions that provided insights into the regulatory interactions within the TRN (Supplementary Information, Text S1).

**Figure 5.**
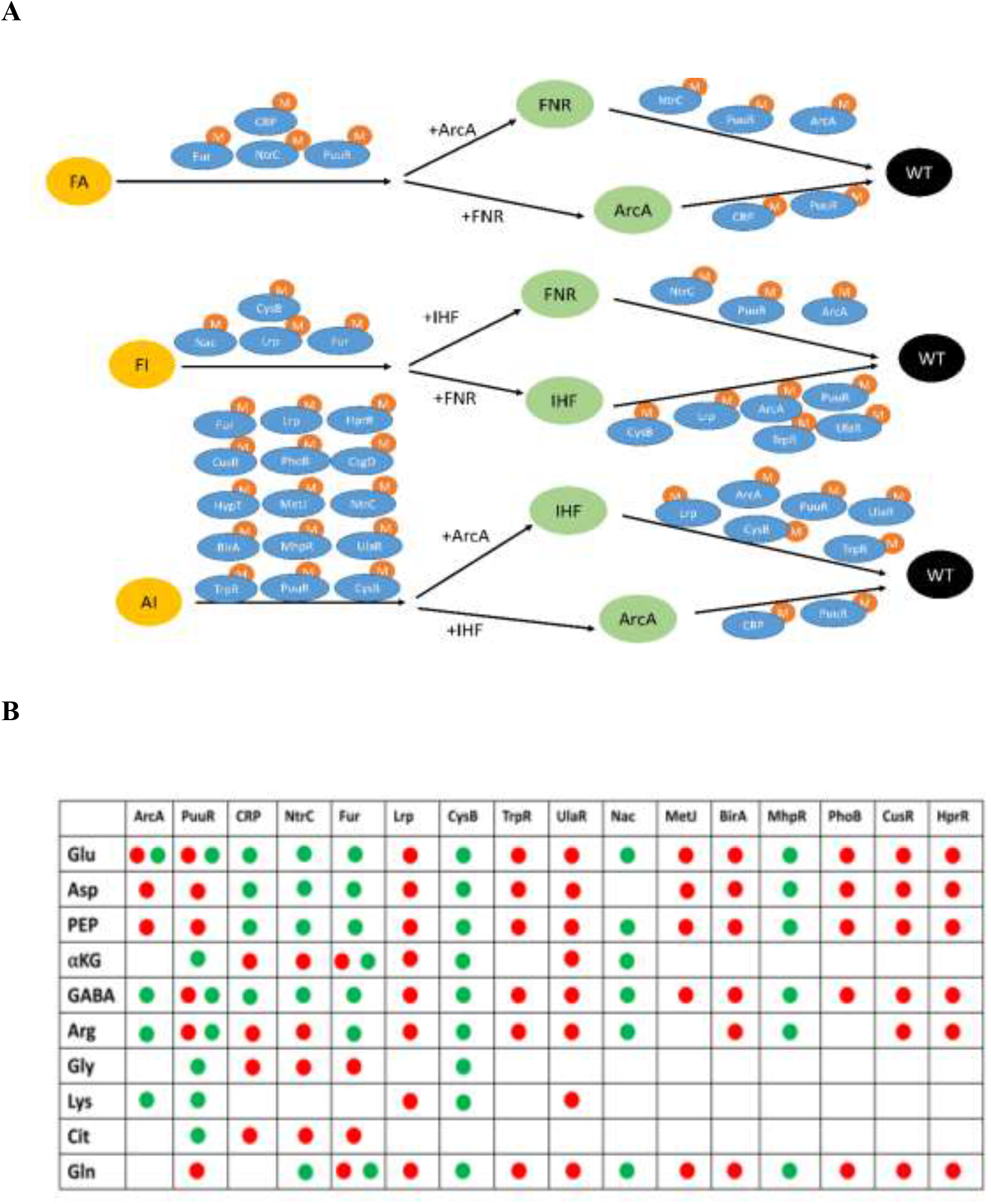
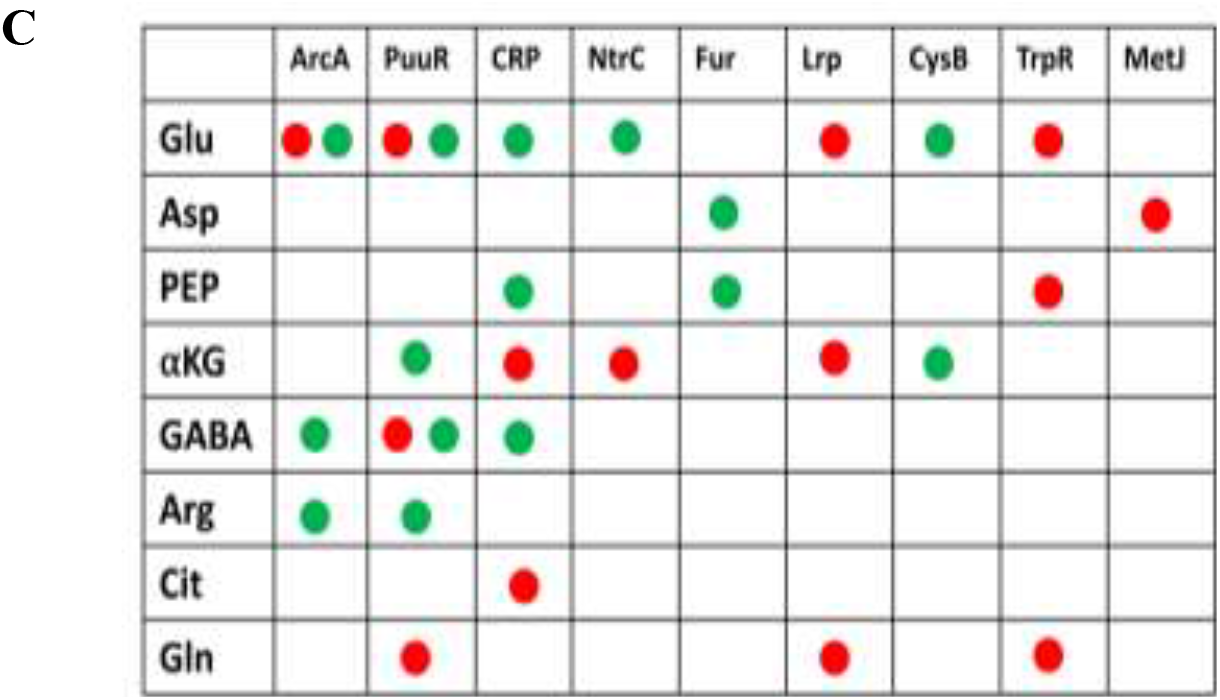
Characterization of conserved and novel metabolite-TF interactions in each of the double and its corresponding single mutants. **A)** “M” represents metabolites (characterized in Figure 5B) that allosterically modulate the iTF activity. The arrangement of the double and single mutants is based on their specific growth rates with the fastest growing WT on the extreme right and slower growing FA on the extreme left. **B)** Comprehensive identification of putative metabolite-iTF interactions using the Pearson correlation (P < 0.05) of metabolite concentrations with iTF activity observed in each of the strains. The green and red dot represents positive and negative correlations respectively. **C)** Subset of putative but high confident metabolite-iTF interactions with Pearson correlation (r > 0.9, P < 0.01) that belong to the same metabolic subsystem where this iTF regulates as well as those putative interactions reported in literature. Abbreviations: Glu, Glutamate; Asp, Aspartate; PEP, Phosphoenol pyruvate; aKG, α-ketoglutarate; GABA, γ-aminobutyric acid; Arg, Arginine; Gly, Glycine; Lys, Lysine; Cit, Citrate; Gln, Glutamine.

### Co-expression analysis of the direct and indirect KPE DEGs

Though a significant proportion (30%) of the indirect or growth mediated genes reflected the metabolite-driven allosteric regulation, a higher proportion (70%) of the genes did not show any significant (Fischer’s exact test, P < 0.05, BH adjusted) enrichment for any iTF binding. These remaining set of genes were annotated to be predominantly regulated by RNA polymerase-associated sigma factors either by using EcoCyc or based on promoter prediction. RNA polymerase-associated sigma factors were used because it is known to be an indirect readout of growth rate-dependent global machinery (Berthoumieux et al., 2013; Gerosa et al., 2013; Klumpp et al., 2009). The genes whose promoters were not annotated to RNA polymerase-associated sigma factors in EcoCyc, were predicted independently using iPromoter-2L (Chen et al., 2014; B. Liu et al., 2018) (Supplementary File S1). However, the occurrence of such indirect genes along with specific direct genes is unclear, which could be better explained if looked through the underlying co-expression patterns. To gain a systematic understanding of the coordinated changes in direct targets and indirect targets including those driven by the underlying metabolite-iTF interaction, we performed weighted gene coexpression network analysis (WGCNA) (Langfelder and Horvath, 2008). WGCNA is an unsupervised technique to cluster genes using correlation of their expression profiles. Such analyses would also reveal the co-regulated genes directly modulated by the dTF/s, which has been a major challenge in understanding the regulatory principles (Imam et al., 2015; Tsai et al., 2018). We employed a large-scale compendium gene expression data (COLOMBUS (Meysman et al., 2014) comprising of microarray and RNA seq data as an input for generating the network of co-expressed genes in *E. coli* (Supplementary Information, Figure S8A and B). Among 4077 contrasts and 4189 genes used for the analysis, we observed occurrences of 7 coexpression merged modules (black, blue, brown, green, red, turquoise and yellow) (Supplementary File S1). Among these modules, blue (1274) and turquoise (1572) covered the majority (~68%) of the genes. Such analyses reflected that the occurrences of particular gene expression were either based on a specific function or co-occurring/co-existing functions. We performed gene ontology (GO) analysis (as described in methods section) to identify the functions of the KPE genes falling under a particular module for both the regulation-specific and growth-rate specific upregulated and downregulated genes in each of the mutants. It should be noted that all the regulatory targets in the growth-rate specific genes were separately enriched for module and GO analysis. By such formalism, we were able to comprehensively illustrate the functions of either regulation-specific or remaining indirectly affected genes modulated by the dTFs and iTFs. Besides, this gave us insights regarding the preference for the metabolite-iTF interaction that modulated a specific set of genes.

We could also identify the coregulated genes and their functions controlled by the regulators FNR, ArcA, and IHF as described by the module colours. Further, the GO categories enriched for each of the modules in the regulation-specific genes were found to be similar to the enrichments seen in the growth mediated genes (Figure 6 and Supplementary Information, S9A-D). This indicated not only the coordination in genes under direct regulatory control that were co-expressed with the growth-rate mediated genes but also the importance of such indirect targets. Indeed, across the mutants, we found either enrichment for generation of precursor metabolites or deoxyribonucleotide metabolism or protein transport in the blue module in the regulation-specific downregulated genes that showed co-expression with negative regulation of macromolecule biosynthesis in the blue module of growth-rate mediated downregulated genes. Such co-expression patterns between the direct and indirect genes underscore the constraints on the precursor or metabolite usage. Similarly, we found either enrichment for α-amino acid metabolic process or arginine biosynthetic process or glutamate metabolic process in the turquoise module in the regulation-specific downregulated genes that showed coexpression with cellular compound nitrogen biosynthetic process in the turquoise module of downregulated indirect targets. Moreover, in the case of the red module of regulation-specific and growth-rate mediated upregulated genes, we found co-expression of genes enriched for TCA cycle or fatty acid metabolism or aerobic respiration with putrescine catabolic process in the majority of the mutants. This could reflect the re-generation of αKG through the putrescine degradation as the upregulation of the aerobic TCA cycle genes can result in its futile cycling. Most of the metabolite-driven TF targets belonged to either red or turquoise modules and their functional enrichments also revealed similar co-expression patterns specific to the modules. Overall, this indicated that if the direct effects of the dTFs resulted in changes in the biosynthetic process (turquoise model), the metabolite-driven iTFs concomitantly regulated alternative biosynthetic genes (Text S1), thereby regulating the economy of the metabolite. This explains the rationale behind the regulatory interactions associated with the deleted regulator and also the set of genes that were modulated. Hence, we were able to dissect the complexity of regulation on gene expression changes using WGCNA.

**Figure 6:**
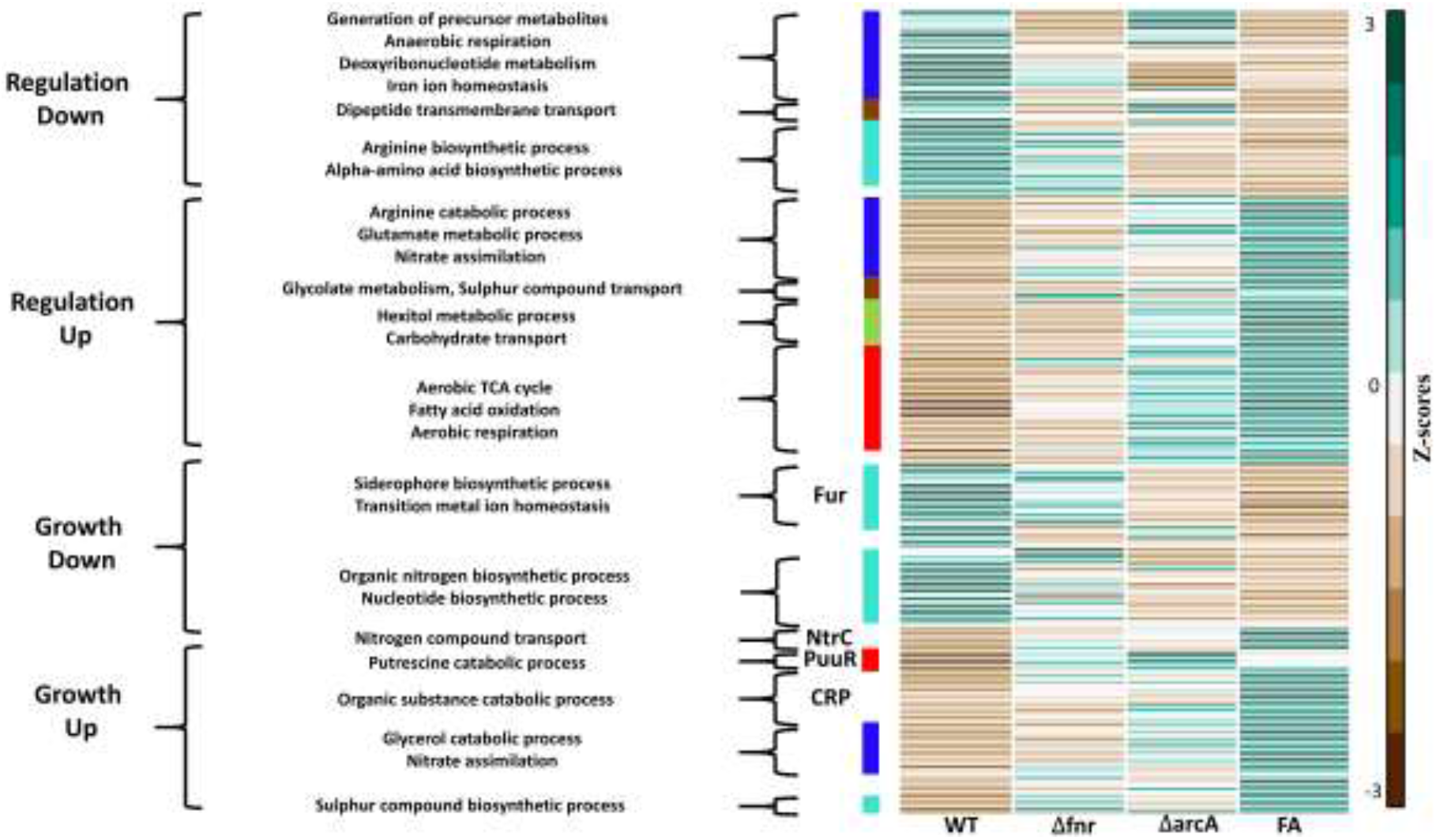
Heatmap depicting the regulation-specific direct and growth-rate mediated indirect targets for FA compared to WT. Both the upregulated and downregulated KPE DEGs in the FA mutant compared to WT were used to compare across the strains. The tpm (transcript per million) values calculated for each of the double mutant and corresponding single mutant strains were z-score transformed. The colour strips next to the heatmap represent the different co-expression modules displayed here as different colours. The GO categories or TFs that have genes belonging to more than 1 module colour are shown as blank regions between the coloured strips. See Figure S9 A-D for FI and AI.

## Discussion

To gain a clear understanding of the regulatory logic, disentangling the direct and the indirect pleiotropic effects that exist within the TRN becomes imperative. This work attempted to decipher the complex layers of TF-driven decision making on glucose fermentative metabolism in *E. coli*. Here, we characterized and quantified the pleiotropic effects on gene expression coordinated by the TFs as well as growth-rate dependent global machinery (RNA polymerase and intracellular metabolites) under exponential growth of the bacteria. Overall, we found across the strains lacking FNR, ArcA, or IHF and their combinations thereof, that a greater percentage (>45%) of DEGs were associated with the direct effects of the knocked-out TF as opposed to earlier estimates (~17%) (Iyer et al., 2021). This was determined from the available TF binding data, novel genes wherein the upstream DNA binding sequence was predicted with good confidence and chemostat based gene expression analyses. These direct targets represent “regulation-specific” genes and are devoid of growth-rate effects. Further, we identified the indirect gene targets in batch fermentative conditions for each of the TF-deleted strains. These indirect targets of the dTFs were determined from the gene sets modulated by growth-rate in terms of RNA polymerase sigma factor distribution as well as the interactions with other iTFs based on metabolite regulation. Our study reinforced the widespread growth rate-mediated effects on gene expression known to be controlled by RNA polymerase, ribosome and intracellular metabolites (Berthoumieux et al., 2013; Gerosa et al., 2013; Klumpp et al., 2009; Scott et al., 2014; Yu et al., 2021). Functionally, both the direct and indirect targets of the dTFs not only represented the metabolic bottleneck genes but also hedging or alternate carbon metabolism genes that will facilitate growth in new environmental conditions. Further, we infer that the distribution of the sigma factors across the indirect genes could also explain their gene expression changes in response to nutrient limitations (majorly carbon and nitrogen and to a lesser extent phosphate or sulfur) as a result of reduced growth rates (Hua et al., 2004; Simen et al., 2017; Yu et al., 2021).

Despite the overriding complexity within the TRN, our findings demonstrated the nature of dTF-iTF interactions and their effect on regulation of gene expression. Except a few, we found that majority of the gene expression changes modulated by the enriched iTFs could be ascribable to their post-translational effects, highlighting the metabolite-driven regulation. Additionally, a simple correlation analysis gave us insights into the specific interactions of the dTF with the dense web of iTFs, by identifying the direct target specific metabolites perturbed by the dTF and determining the activity of iTFs using NCA. Among the total putative metabolite-iTF interactions identified with high confidence in this study, we not only validated the metabolite-iTF interactions (~45%) by comparing with published experimental data but also predicted many novel potential metabolite-iTF interactions (~55%). Further, we were able to trace the exact nature and cause of occurrence of specific metabolite-TF interactions using pathway-based analysis. Additionally, we revealed the independent yet additive mode of interactions between the global regulators FNR, ArcA, and IHF using combinatorial deletion of TFs. Overall, this study employing combinatorial deletion approach provides a paradigm for identifying the conserved regulatory pattern across the mutants, the amount of overlap in genes and functions, and target expression specificity arising out of the mode of interactions of the TFs.

Apart from metabolite based dTF-iTF interactions, we considered that indirect gene targets that could not be assigned to known TFs were regulated primarily by RNA polymerase sigma factor. However, it is of particular importance to understand the association between the indirect and direct targets which influence the growth physiology. Towards this, we performed co-expression analysis using WGCNA which revealed the complementary nature of indirect gene targets with direct gene targets in accordance with the metabolite adjustments. For instance, to mitigate the competition for available cellular resources such as precursors and proteinogenic amino acids, if the pathways synthesizing the metabolite is perturbed due to the deletion of the TF, the organism tries to either reduce the usage of the particular metabolite in other pathways or promotes its degradation if found to exceed the demand for the metabolite. Moreover, the co-expressed targets of the deleted TF elucidate the preference for interaction with specific iTFs as well as their downstream cognate genes. Broadly, these co-expressed metabolic genes can be considered as co-regulated genes, given the similar expression profiles and upstream consensus motif sequences for any TF. Overall, the insights obtained from the gene co-expression patterns for *E. coli* K12 (COLOMBUS database) can now be employed to explore gene regulatory mechanisms that enable the organism to cope with stress or changes in carbon sources.

Our previous work have shown the allocation of necessary and unnecessary proteome sector and metabolite adjustments, driven by the global transcriptional factors FNR, ArcA, and IHF (Iyer et al., 2021). Here, we have determined plausible regulatory interactions and crosstalk effects which facilitate favourable cellular phenotype. Hence, using an integrated flow of analysis (Figure 1), we elucidated the multi-level cellular decision-making of the organism in formulating an appropriate biological response for a particular environment. However, we note that the correlation analysis just explores the possibility of occurrence of putative metabolite-TF interactions and does not imply direct causation. Thus, additional work involving the experimental validation of metabolite-TF interactions (Diether et al., 2019; Lempp et al., 2019) as well as characterizing other modes of regulation such as sRNA (Wang et al., 2015) or protein-protein interaction (Arifuzzaman et al., 2006) would further contribute to our comprehensive understanding. Besides, the demarcation of the indirect and direct targets of global TFs are condition-dependent, meaning these indirect gene targets have binding sites for the TF but represent silent targets for that particular environment (Choudhary et al., 2020; Park et al., 2013). These sites could also represent relaxed specificity sites that could enable adaptation to newer environments (Taylor et al., 2022). Additional experiments supporting the upstream binding of the TFs to uncharacterized or putative direct targets will further strengthen our predictions. Nevertheless, this study serves as a valuable approach to accurately deconvolve the complex layers of TF-driven regulation and systems-level regulation, in general.

## Materials and methods

### Strain construction

The *E. coli* K12 MG1655 was used to construct the double deletion strains of Δ*fnr*Δ*arcA* (FA), Δ*fnr*Δ*ihf* (FI), and Δ*arcA*Δ*ihf* (AI). Δ*ihf* represents a deletion of both the subunits *ihfA* and *ihfB* (Supplementary Information, Table S2). The strains were generated using λ-red mediated homologous recombination using the plasmids pKD46, pKD13 as template plasmid, and pCP20 for removal of marker (Datsenko and Wanner, 2000). The gene deletions were verified by PCR as well as Sanger sequencing using primers against the gene of interest. After verifications, stocks were prepared in 25% glycerol (final concentration) and stored at −80°C.

### Media and strain cultivation

The cells were cultivated anaerobically in a 500 ml bioreactor (Applikon) containing 400 ml of M9 minimal media (6 g/liter anhydrous Na2HPO4, 3 g/liter KH2PO4, 1g/liter NH4Cl, 0.5 g/liter NaCl plus 2 mM MgSO4 plus 0.1 mM CaCl2) with 2 g/liter glucose as the carbon source. The cultivations and bioreactor settings were done as reported previously (Iyer et al., 2021). Briefly, the cells were used to inoculate the 400 ml M9 minimal media with 2 g/litre glucose in a bioreactor and the start optical density (OD) of the culture was set at approx. 0.07 OD. The temperature in the bioreactor was maintained at 37°C, stirrer speed at 150 rpm, and pH was maintained at 7.2. The bioreactor was sparged with nitrogen to maintain the dissolved oxygen levels at zero at all times. The slope of the linear regression line fit to the natural logarithm of the optical density values at 600 nm wavelength (OD600) versus time plot, ln (OD600) versus time (in hours) was used to calculate the exponential growth rate of the organism. All the physiological and multi-omics characterizations were done in the exponential phase of the organism. The dry cell weight (DCW) for each strain was experimentally derived during its exponential growth phase wherein 1.0 OD at 600 nm corresponds to 0.44 g DCW h^-1^.

All phenotypic characterizations were performed in three biological replicates (n = 3) as described previously (Iyer et al., 2021). Extracellular supernatants collected from the entire exponential growth phase were used to determine the rates of glucose uptake as well as the rates of secretion of extracellular mixed acid fermentation metabolites such as acetate, lactate, pyruvate, succinate, formate, and ethanol, and the yields thereafter. All physiological characterizations were tested for significance by using an unpaired two-tailed Student’s t-test.

Cell cultivations were performed in a chemostat (Pal et al., 2022; Utrilla et al., 2016),wherein the feed media composition was the same as the batch media. The WT strain was grown under anaerobic batch growth until 80% of the maximum biomass was produced (biomass formation was monitored by OD measurements) or until 80% of the glucose present in the media was consumed (glucose levels were monitored by HPLC) after which addition of the feed media was started. The dilution rate in the chemostat was maintained at 0.21 h^-1^. Once the cells reached the steady-state, the chemostat cultivation was continued for 3-5 residence times after which the cells were harvested for RNA extraction and sequencing. The RNA sequencing data for aerobic batch growth performed under identical chemostat conditions were obtained from (Pal et al., 2022).

### RNA extraction and mRNA enrichment

The RNA extraction for each strain was harvested from the mid-exponential phase for two biological replicates as described previously (Iyer et al., 2021). Library preparation for RNA sequencing was done following a paired-end strand-specific protocol using NEBNext Ultra Directional RNA library kit. RNA sequencing was carried out on HiSeq 4000 rapid run mode using 2×150 bp format at Genotypic Technologies, Bangalore, India. For chemostat cultivations (aerobic and anaerobic), the harvested RNA was extracted and sent for RNA sequencing to C-CAMP NCBS Bangalore, with NEBNext Ultra II Directional RNA library kit (single strand-specific) and run on HiSeq 2500 (Rapid run mode) using 1×50 bp format.

### Transcriptome data analysis

The raw RNA sequencing files from the mid-exponential phase of batch growth and the chemostat cultivations were first trimmed with CUTADAPT (Martin, 2011) to eliminate the adapter sequences and the low-quality reads. The trimmed reads were mapped to the *E. coli* K12 MG1655 genome (GenBank accession number NC_00913.3) using BWA (Li and Durbin, 2009) aligner and sam file was then converted to bam file using Samtools (Li et al., 2009). Counts at each gene level were assigned with FeatureCounts (Liao et al., 2014) using the reference genome provided in GTF format. The 4466 genes used for the analysis were obtained from EcoCyc database (v. 21.5) (Keseler et al., 2017). All rRNA, tRNA, and sRNA genes were excluded from the analysis. Differentially expressed genes were analysed using EdgeR (Robinson et al., 2009) after eliminating the genes having less than 10 reads. The genes with greater than or more increase or decrease in gene expression and had BH-adjusted P <0.05 value were considered to be DEGs and were used for subsequent analysis. The DEGs were then enriched for metabolic pathways using KEGG pathway classification and the significance of each pathway was checked using a hypergeometric test (P <0.05, BH adjusted) in R. The criteria for a pathway to be considered significant was atleast 9 upregulated or downregulated genes. DEGs without any Accession ID (EcoCyc v 21.5) such as phantom genes were not included in the analysis.

In the comparison of transcriptomes between the WT anaerobic fermentation versus the WT aerobic respiration from glucose-limited chemostat cultivations, the DEGs were annotated as ‘Regulation-specific’ (electronic supplementary material, file S1) and represent genes that are not altered due to changes in growth rate or slow growth rate. These DEGs are filtered with the threshold of (Benjamini-Hochberg adjusted) P less than 0.05 and at least 2-fold change. These genes were then used to identify the Regulation-specific and growth rate mediated genes in the transcriptome changes of single or double mutants compared to WT in batch mid-exponential phase. These regulation-specific genes were further validated using previously reported direct gene targets of these regulators from *in vivo* ChIP binding data as well as binding motif prediction analysis using FIMO from the MEME suite (Charles E. Grant et al., 2011). The global and pathway-specific local regulators together with their cognate gene targets were obtained from RegulonDB (v 10.6.3) (Santos-Zavaleta et al., 2019). The KPE DEGs were independently enriched for global and local TFs and significance was assessed by Fischer’s exact test (BH-adjusted, P < 0.05) with an arbitrary set of 5 genes. It should be noted that even though the deleted TF can be enriched in the case of indirect gene sets, they were not considered for analysis based on the assumption that these genes have other TF binding sites that overlap or simply they are bound yet represent silent targets (Choudhary et al., 2020; Park et al., 2013) for this particular condition. Further, enriched TFs which have dual regulation were considered neither for NCA nor for correlation analysis.

Gene ontology analysis was performed using the AmiGO 2 database (Carbon et al., 2009) or topGO package (Alexa and Rahnenfuhrer, 2020). The top categories with either FDR < 0.05 or P < 0.05 that accounts for a larger set of genes were considered for further analysis.

### Metabolomics

#### 1. Extraction of metabolites

The metabolites extraction for each of the strains was performed in the mid-exponential phase. Metabolite extraction was performed for 3 biological and 2 technical replicates (n = 6). A fast-cooling method was employed to quench the harvested cells as described previously (Iyer et al., 2021; Pal et al., 2022). ^13^C labelled extracts were used as internal standards generated by growing WT *E. coli* K12 MG1655 cells aerobically and were then used to quantify the key metabolite pool sizes using an isotope-based dilution method as done previously. During the earliest stage of extraction, each of the samples was spiked with a fixed volume of internal standard. The volume of the pooled internal standard was added to each sample such that the external and internal standard peak heights differed less than 5-fold. Following the sample preparation steps (Iyer et al., 2021; Pal et al., 2022), the samples were completely dried in a vacuum concentrator and stored at −80°C. Before analysis in an LC-MS/MS instrument, samples were reconstituted in 100 μl of chilled (−20°C) acetonitrile: buffered water (60:40, v/v) and centrifuged at 4°C for 10 mins and the supernatant was transferred to prechilled glass vials. Buffered water was composed of 10 mM ammonium acetate (pH 9.23). pH was adjusted using ammonium hydroxide prepared in HPLC-grade water.

#### 2. LC-MS/MS settings

The samples for metabolomic analysis were run on a high-resolution mass spectrophotometer in Orbitrap Q Exactive Plus (Thermo) equipped with a SeQuant ZIC-pHILIC column (Merck) of dimensions 150 mm x 2.1 mm x 5-micron packing and in sequence with a ZIC-pHILIC guard column (Merck) of dimensions 20 mm x 2.1 mm x 5-micron packing with the LC-MS/MS settings and calibrations as described previously (Iyer et al., 2021; Pal et al., 2022). An alkaline mobile phase was used with electron spray ionization (ESI) ion source operated in positive (M + H)+ and negative (M - H)-mode separately. A full scan range of 66.7 to 1000 m/z was applied for positive as well as negative modes and the spectrum data type was set to profile mode. The mobile phase used for chromatographic separation consisted of a nonpolar phase A (acetonitrile: water mixed in the ratio 9:1, 10 mM ammonium acetate, pH 9.23 using ammonium hydroxide) and polar phase B (acetonitrile: water mixed in the ratio 1:9, 10 mM ammonium acetate, pH 9.23 ammonium hydroxide). A linear gradient with flow rate of 200 μl/min was set as follows: 0–1 min: 0% B, 1–32 min: 77.5% B, 32–36 min: 77.5% B to 100% B, 36–40 min: hold at 100% B, 40–50 min: 100% B to 0% B, 50–65 min: re-equilibration with 0% B. An injection volume of 5 μl was used for all the samples and standards.

#### 3. Metabolomics data analysis

The raw data from the machine was assessed using the Xcalibur 4.3 (Thermo Fisher Scientific) Quan Browser software package as done previously (Iyer et al., 2021; Pal et al., 2022). A semi-quantitative analysis was performed using peak heights of precursor ions with a signal/noise (S/N) ratio of more than 3, a retention time window of less than 60 s, and less than 5 ppm mass error. Height ratios of metabolites were obtained after normalizing the peak heights of the samples to the peak heights of the internal standards used. The absolute concentrations of metabolites were determined using ^12^C -labelled chemical standards (mix of 40 metabolites) as described previously (Iyer et al., 2021; Pal et al., 2022). Metabolite concentrations were imputed for missing values and biomass normalized. These values were g-log transformed for identification of the statistically significant metabolites using MetaboAnalyst. Metabolites with false-discovery rate (FDR) < 0.05 (2-tailed unpaired Student’s t-test) were considered for further analysis. The absolute metabolite concentrations were expressed as micromoles per gram dry cell weight (μmol/gDCW) or (Height ratio/gDCW).

### Network component analysis (NCA)

NCA was performed using the MATLAB scripts and codes made available at github from reference (Lempp et al., 2019) with minor modifications. Only the network component analysis and bootstrap code files were used in this study. NCA works on determining TF-activity by least square optimization between the log10 transformed gene expression data (in TPMs) and product of connectivity (a priori known interactions between the TF and cognate genes) and TF-activity (Liao et al., 2003). NCA was performed on a smaller dataset especially on the growth-rate specific or indirect gene sets that belonged to the KPE upregulated and downregulated DEGs. As we used a smaller dataset, only regulators which were able to reproduce the activity profiles similar to manual interpretation were retained for the analysis. For instance, if a TF positively regulates 4 to 5 genes and majorly is an activator based on the a priori knowledge and if this TF is showing activity values of a repressor or inconclusive, then the TF is excluded from further analyses. We were able to successfully reproduce the TF interactions in more than 80% of the TFs across all the growth-rate specific genes in the up and downregulated KPE DEGs for all the strains. An RNA polymerase that connected all the genes within a set was retained. Analysis of sigma factors such as sigma 70 or 38 along with ppGpp was excluded. The simulation was performed in triplicates for each condition along with bootstrapping to ensure confidence in smaller datasets (Misra and Sriram, 2013) as well as to keep the data symmetric for correlation with metabolite concentrations.

### Metabolite-TF interactions

We analyzed the target enrichments of iTFs available from RegulonDB and EcoCyc in the KPE DEGs restricted to indirect gene targets. The activity of transcriptional regulators was determined based on the mode of interaction with its target genes. For instance, if the TF is a repressor and the gene targets of the regulator showed a reduction in gene expression, then we inferred that the activity of the TF has increased. A similar analogy was applied in the case of the activator. However, if the TF had less than 70% of its targets as positively or negatively regulated, then the regulator was assigned to be a dual regulator. TFs whose interaction with their complete gene lists was not conclusive (< 70%) were excluded from the correlation analysis. For instance, if a TF regulates 10 genes of which gene expression of 7 are repressed and 3 are activated then such TF is majorly regarded as a repressor and retained for analysis. However, if expressions of 5 genes were activated and the rest 5 genes were repressed, then the TF is supposedly functioning as a dual regulator and such regulator/s were excluded from the correlation analysis. Further, the TFs whose activities were not reproducible in NCA in terms of their manual interpretation were also excluded from the correlation analysis. Next, the metabolites that fall under the direct targets of the dTFs were retained for correlation analysis. The logic is that the deletion of TF results in changes in gene expression of the direct targets followed by metabolite changes associated with these genes and these metabolites affect allosterically the activity of other iTFs that regulate part of the growth rate mediated or indirect gene sets. To identify the direct target specific metabolites, we performed joint pathway analysis (Metaboanalyst (Xia et al., 2009)) using both of the upregulated and downregulated direct targets/genes with statistically significant metabolite changes for each of the mutants. The pair (atleast 1 metabolite with many genes or 1 gene with many metabolites) which had the impact > 0.1 (Metaboanalyst) were retained for the correlation analysis. The value for the cutoff was arbitrarily set. A Pearson correlation analysis were performed to identify the metabolite-TF interactions. The correlation analysis should be looked with caution as it only explores the possibility of occurrence of metabolite-TF interaction and are not confirmatory in nature. An initial screening of metabolite-TF interactions involved selecting metabolite-TF pairs that have a Pearson correlation coefficient > 0.75 but statistically significant (P < 0.05). Later, those with correlation coefficient, r >0.9 and statistically significant (P < 0.01) or those obtained from literature but found to be significant (r > 0.8, P < 0.05) in our dataset were retained for further analysis.

### WGCNA analysis on Compendium gene expression data

The compendium gene expression data for *E. coli* K12 was obtained from COLOMBUS database (Meysman et al., 2014). COLOMBUS has gene expression data from microarray and RNA seq experiments for various environmental conditions and is further normalized to make it comparable across these conditions. To enable the data suitable for co-expression network analysis, the data without centering or scaling was used to impute missing values for the large dataset using the global SVD impute function (Liew et al., 2011; Troyanskaya et al., 2001). It was performed using pcaMethods (Stacklies et al., 2007) with parameters (no of pc components = 5, threshold = 0.01, maxSteps = 1000, cross-validation cv = “q2”). Further, the expression data was assessed by goodSamplegenes function from WGCNA (Langfelder and Horvath, 2008) package in R. Finally, the total number of genes and samples considered for each organism was: 4189 genes and 4077 samples. For construction of co-expression networks, the scale-free topology was computed using the pickSoftThreshold function and biweight midcorrelation with default parameters to obtain robust modules from a larger dataset. This was followed by construction of signed topology overlap matrix (TOM). The co-expressed genes were clustered into modules using the flashClust function with minimum module size set to 30. The cuttreeDynamic function was used to merge the highly similar modules based on eigengenes and correlation value of 0.75. Each module was represented using a color and if not mentioned otherwise, the gray color corresponds to uncorrelated genes. In the figures and for all the downstream analysis, the genes that belong to a module color and are regulated by iTF (whether enriched or not) are shown/categorized separately.

### Regulator prediction for promoters of direct gene targets

The potential direct targets obtained from chemostat analysis and which did not have reported literature (EcoCyc) or experimental backing were re-analyzed using the FIMO tool (5.4.1) from the MEME suite. The consensus motif used for the analysis for Fnr (5’-TTGATNNNNATCAA-3’ or 5’-TTGATYWNNATCAA-3’), ArcA (5’-GTTAATTAAATGTTA-3’ or 5’-GTTAAWWAAATGTTA-3’), and IHF (5’-DTWYYYNGYNNATTTW-3’ or 5’-WWWNWWNWTNTTW-3’) regulators was obtained from the literature (Federowicz et al., 2014; Grainger et al., 2007; Keseler et al., 2017; Myers et al., 2013; Park et al., 2013; Prieto et al., 2012; Santos-Zavaleta et al., 2019). Statistically significant motifs (P < 0.01) were retained such that the binding sites were strictly within 250 bp upstream of the transcription start site of the target gene. Any motif found within the coding region of any gene was excluded from the analysis.

### Sigma factor prediction for promoters of indirect gene targets

For indirect genes wherein no specific global or local regulator could be assigned by either databases or despite having binding sites the regulators were not significantly enriched, such genes were considered to be just under the control of RNA polymerase sigma factors. Promoter prediction was done either using iPromoter2L (Chen et al., 2014; B. Liu et al., 2018) for genes that did not have an annotated RNA polymerase sigma binding site or predictions from EcoCyc. An upstream 200 bp from the start codon of the gene was chosen for the analysis. The region size (200-81 bp) was accordingly adjusted such that they do not overlap with any coding or promoter regions of other genes. In case of absence of annotation to any of the sigma factors or if genes were arranged in an operon, then the region upstream of the first gene or the nearest gene was used for the sigma factor identification. The choice and order of arrangement of the predicted sigma factors was done based on the frequency and closeness to the transcriptional start site.

## Data deposition

The RNA sequencing data and the processed files from this study are available at NCBI Geo (https://www.ncbi.nlm.nih.gov/geo/query/acc.cgi?acc) with accession GSE195954. The metabolomics data presented in this study is available at the NIH Common Fund’s National Metabolomics Data Repository (NMDR) website, the Metabolomics Workbench, https://www.metabolomicsworkbench.org where it has been assigned Project ID PR001316.

## Acknowledgments

This work was supported by DST fellowship/grant (SB/S3/CE/080/2015) and DBT fellowship/grant (BT/PF13713/BBE/117/83/2015) awarded to K.V.V., and DST-WOSA grant (grant number: SR/WOS-A/ET-58/2017) awarded to Dr. Sumana Srinivasan.

M.S.I. acknowledges Department of Biotechnology (DBT), Government of India for his fellowship (DBT/IIT-P/323). A.P. acknowledges Department of Science and Technology INSPIRE (DST-INSPIRE), Government of India for her fellowship (IF140914).

We thank Dr. Sumana Srinivasan for her valuable inputs and funding for the completion of the study. We acknowledge Genotypic Technologies, Bangalore for library preparation and RNA sequencing. We thank Dr. Mayuri Gandhi (Research Scientist, SAIF, IIT Bombay) for providing access to the central LCMS facility. We thank Prof. Pradeepkumar P. I. (IIT Bombay) for providing RT-PCR facility and vacuum concentrator for our use.

## Author contributions

M.S.I. and K.V.V. designed research; M.S.I. and A.P. performed experiments, analysed data and performed computational modelling; M.S.I. and A.P. wrote the paper. K.V.V provided edits, comments and revised the paper. All authors agreed to the final manuscript.

