## Supplementary information for "A systems biology approach to disentangle the direct and indirect effects of global transcription factors on gene expression in *Escherichia coli*"

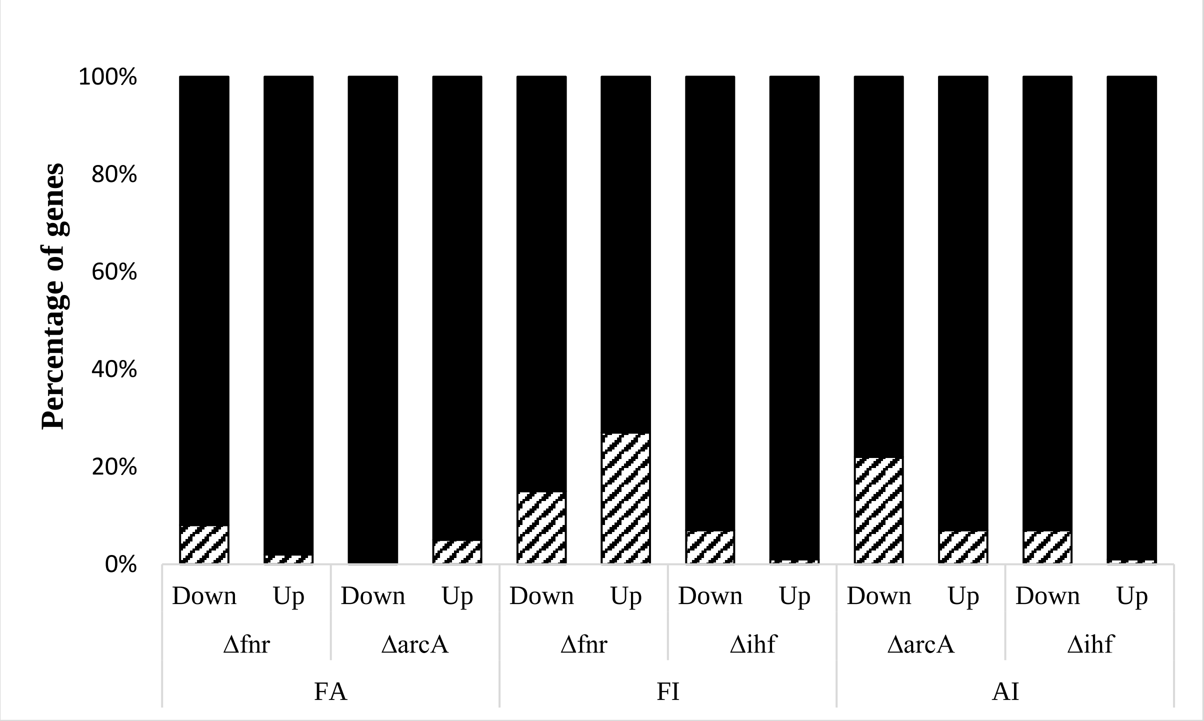


**Figure S1.** Stacked plot depicting the percentage of genes in the double mutants that followed similar direction (log2FC < 0 or > 0) of gene expression as in downregulated and upregulated DEGs in the single mutants. The black region represents the genes that followed the same direction of expression in case of the double mutants and were DEGs in the single mutants. The checkered region represents the genes that did not follow the direction of gene expression.

**A**

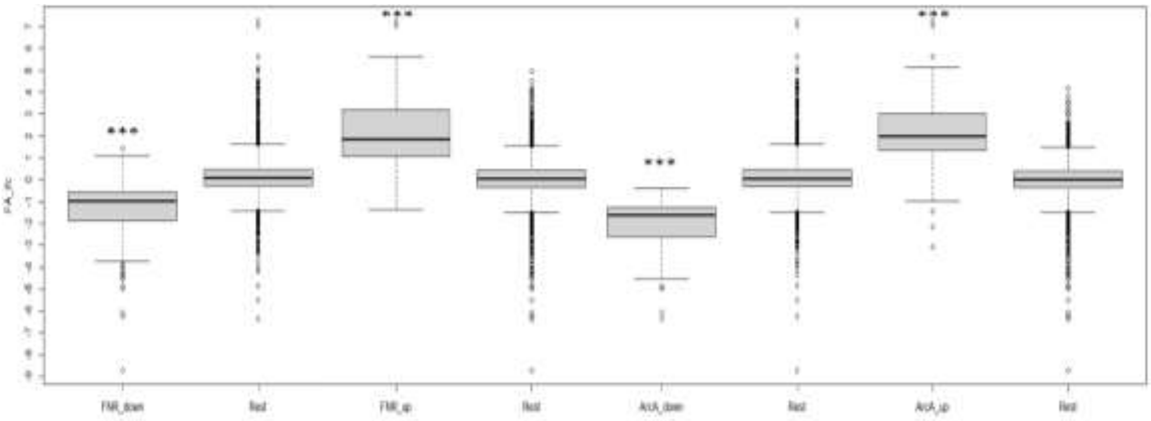


**B**

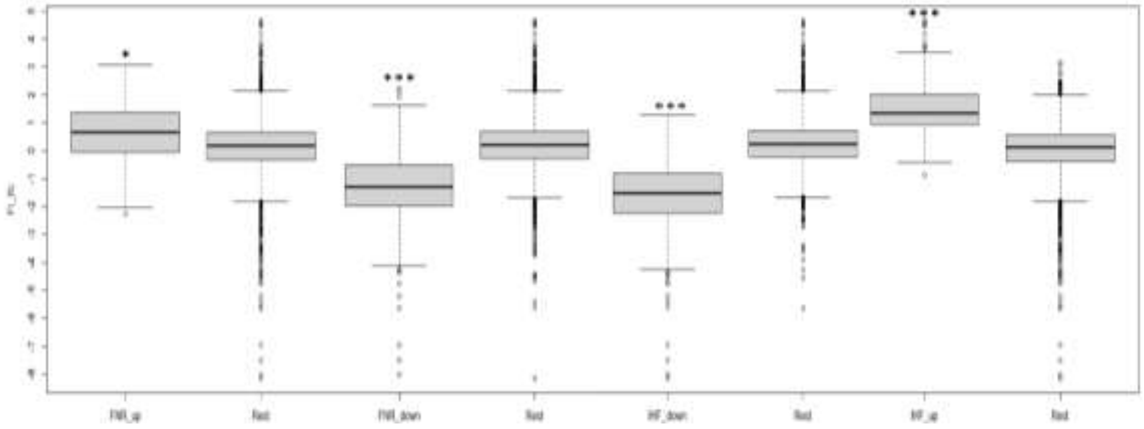


**C**

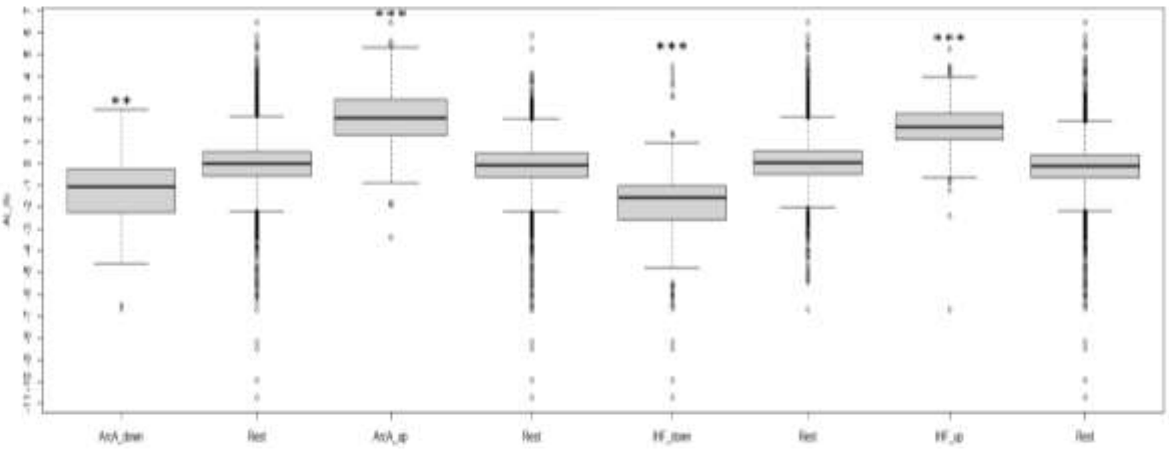


**Figure S2.** Boxplots of log2 fold change in gene expression in **A)** FA, **B)** FI and **C)** AI strains (compared to WT) which was significantly downregulated or upregulated in the single mutants compared to rest of the genes. Significance is represented by asterisks wherein * indicates $P < 10^{-7}$, ** indicates $P < 10^{-12}$ and *** indicates $P < 10^{-15}$ (Wilcoxon test).

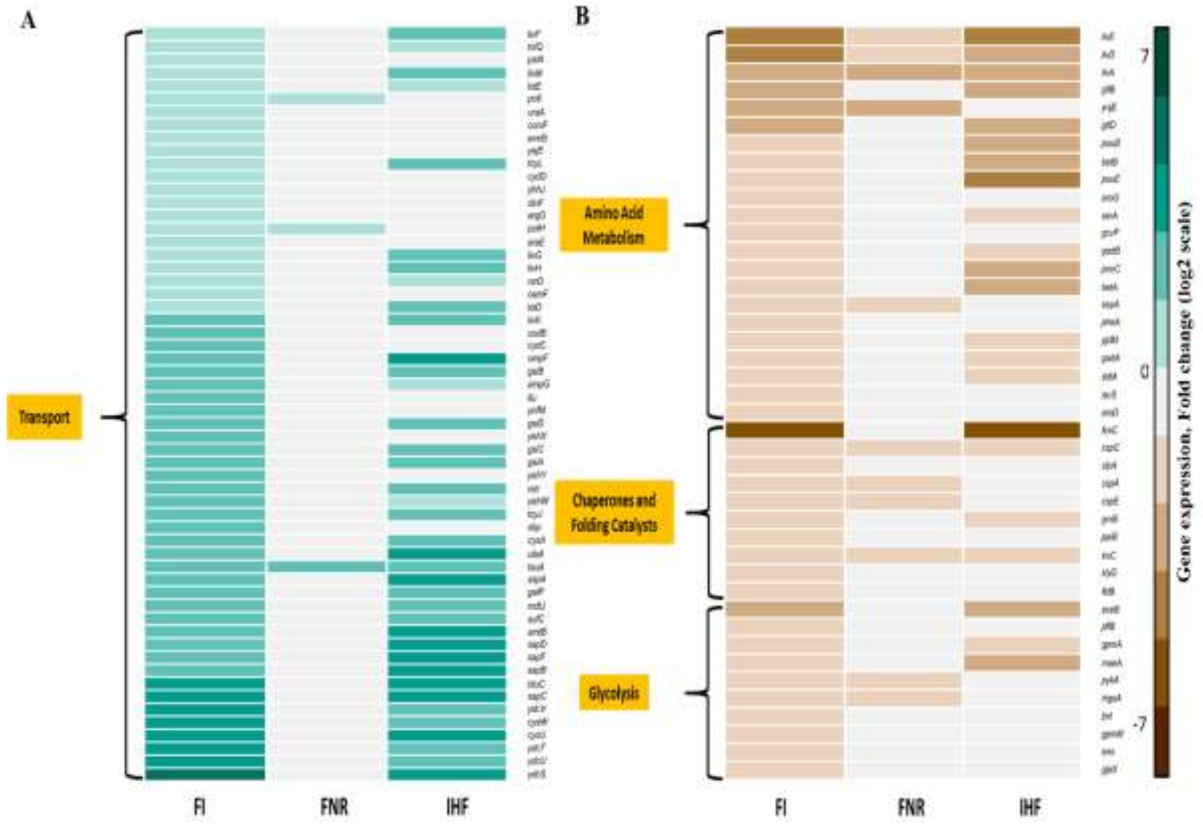


**Figure S3.** Heatmap depicting the metabolic pathway analysis of regulator mutants compared to WT. The figure shows the comparison of A) upregulated and B) downregulated DEGs of FI mutant compared to WT. Only the significantly enriched KEGG pathways are shown. For comparison across the strains, only the DEGs found in FI, Δ*fnr* and Δ*ihf* mutant compared to WT were considered.

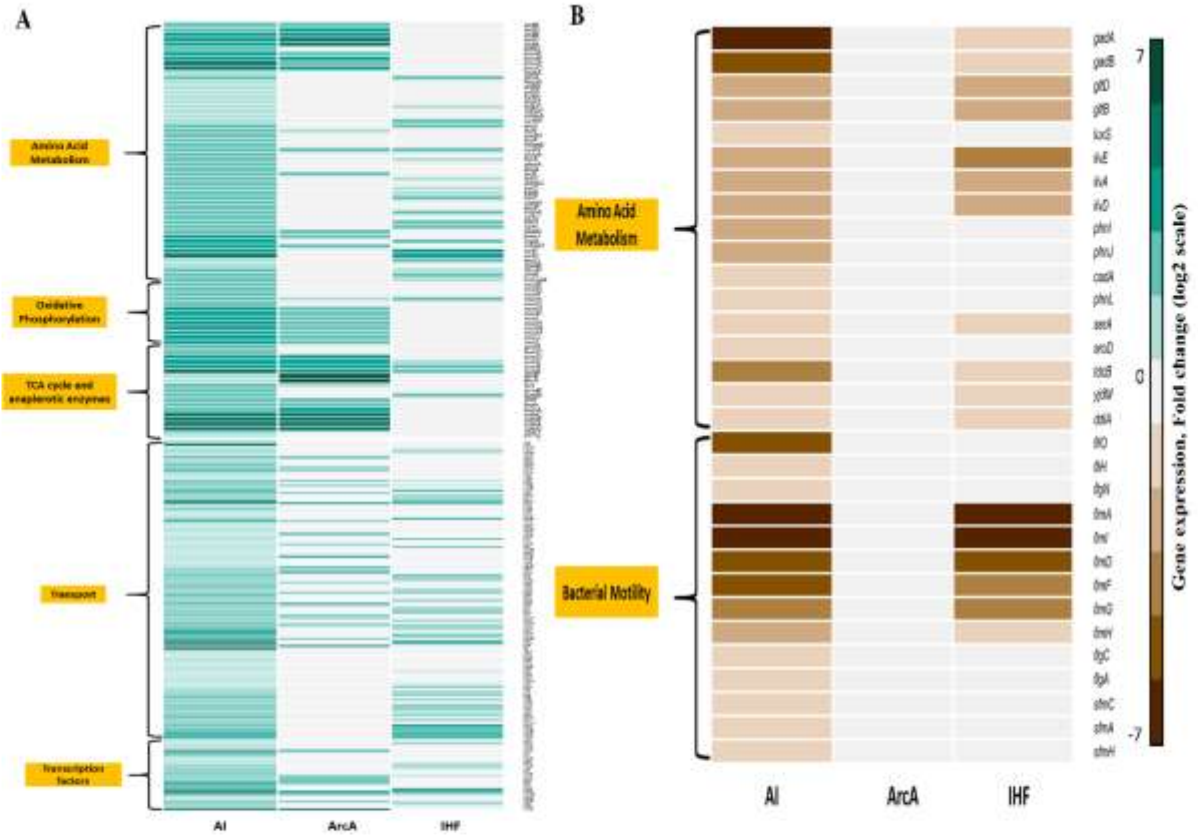


**Figure S4.** Heatmap depicting the metabolic pathway analysis of regulator mutants compared to WT. The figure shows the comparison of A) upregulated and B) downregulated DEGs of AI mutant compared to WT. Only the significantly enriched KEGG pathways are shown. For comparison across the strains, only the DEGs found in AI, Δ*arcA* and Δ*ihf* mutant compared to WT were considered.

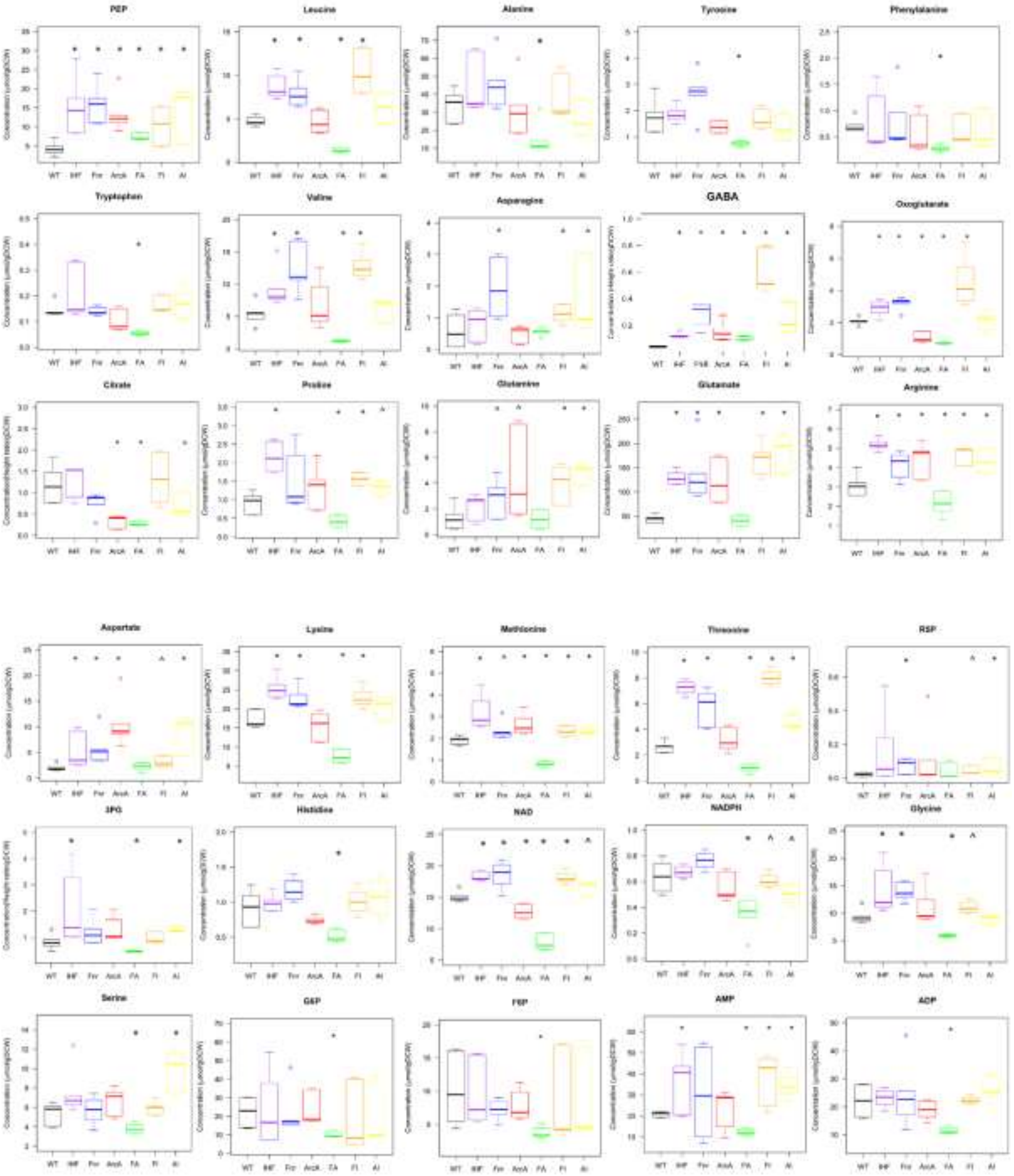


**Figure S5.** Metabolomic analysis of central carbon metabolites perturbed as a result of deletions of global TFs FNR, ArcA and IHF in single and double combinations. Only the significantly (FDR < 0.05, asterisk) altered metabolites in at least one of the mutants are displayed as boxplots. For comparison across the strains, metabolite levels whose significance exceeds FDR = 0.1 (shown as caret) are excluded.

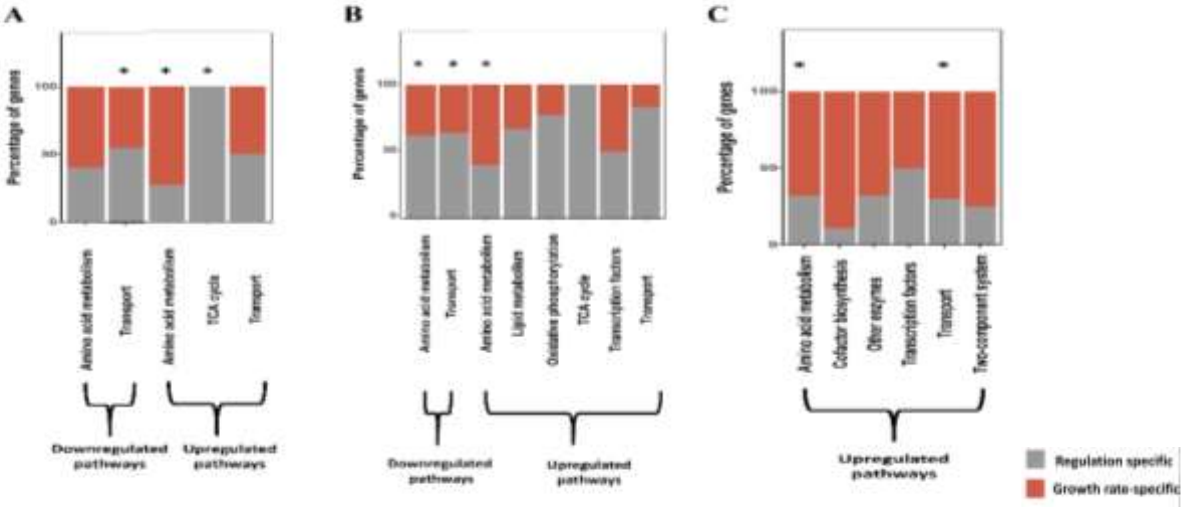


**Figure S6.** Direct and indirect effects on gene expression. Stacked plot depicting the percentage of direct targets and growth-rate mediated indirect targets in **A)** Δ*fnr*, **B)** Δ*arcA*, **C)** Δ*ihf* mutants compared to WT. Asterisk represent the significantly enriched metabolic pathways. Only the metabolic pathways which had atleast 9 genes were retained for the analysis. See Supplementary File S1 for the complete list.

**A**

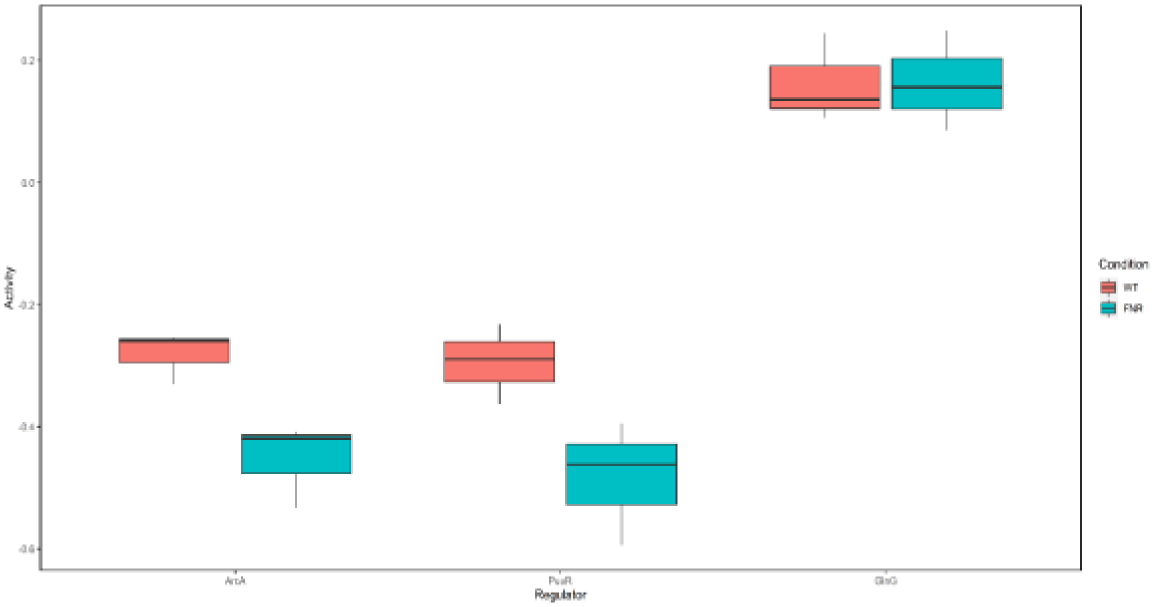
Activity
Regulator
Condition
WT
FNR

**B**

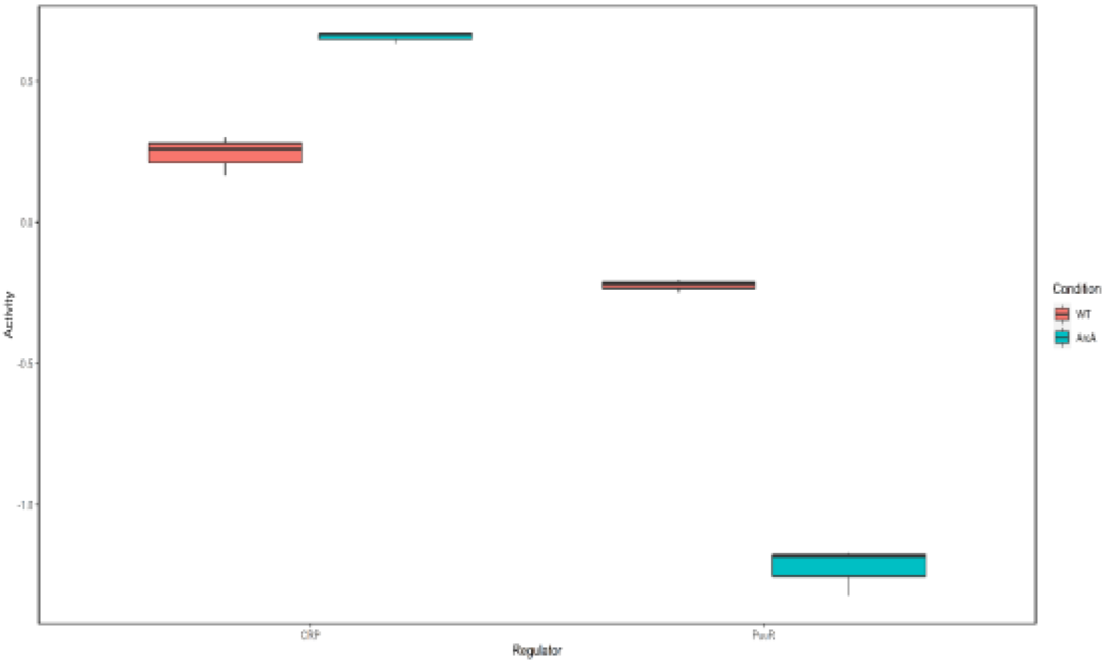
Activity
CRP
Regulator
Condition
WT

**C**

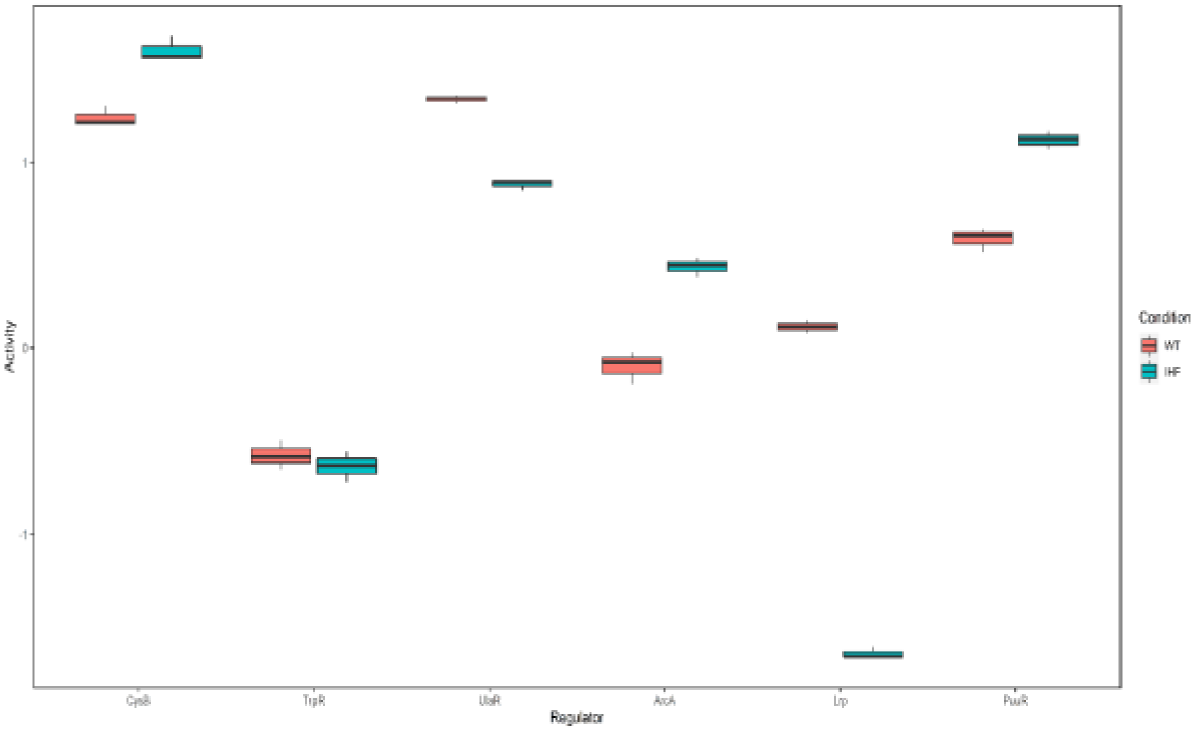
Activity
1
0
-1
CysB
TrpR
UlaR
ArcA
Lrp
PuuR
Regulator
Condition
WT
IHF

**D**

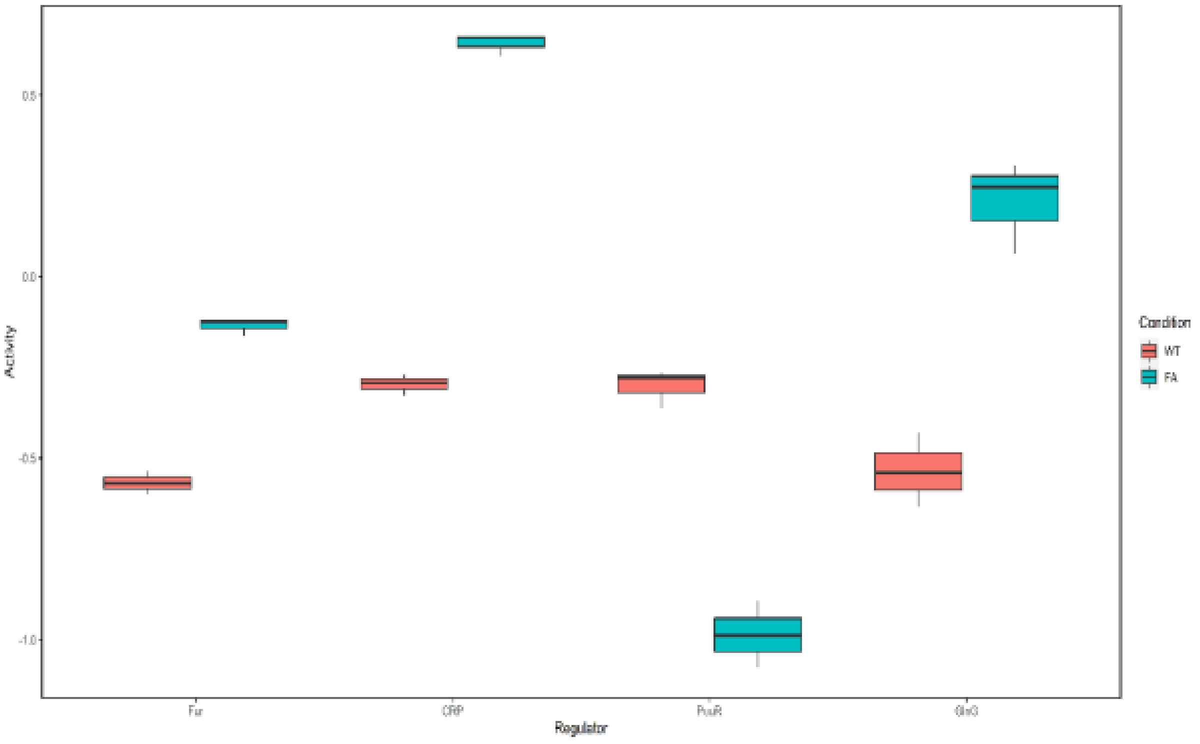
Activity
0.5
0.0
-0.5
-1.0
Fur
CRP
PuuR
Regulator
Condition
WT
FA

**E**

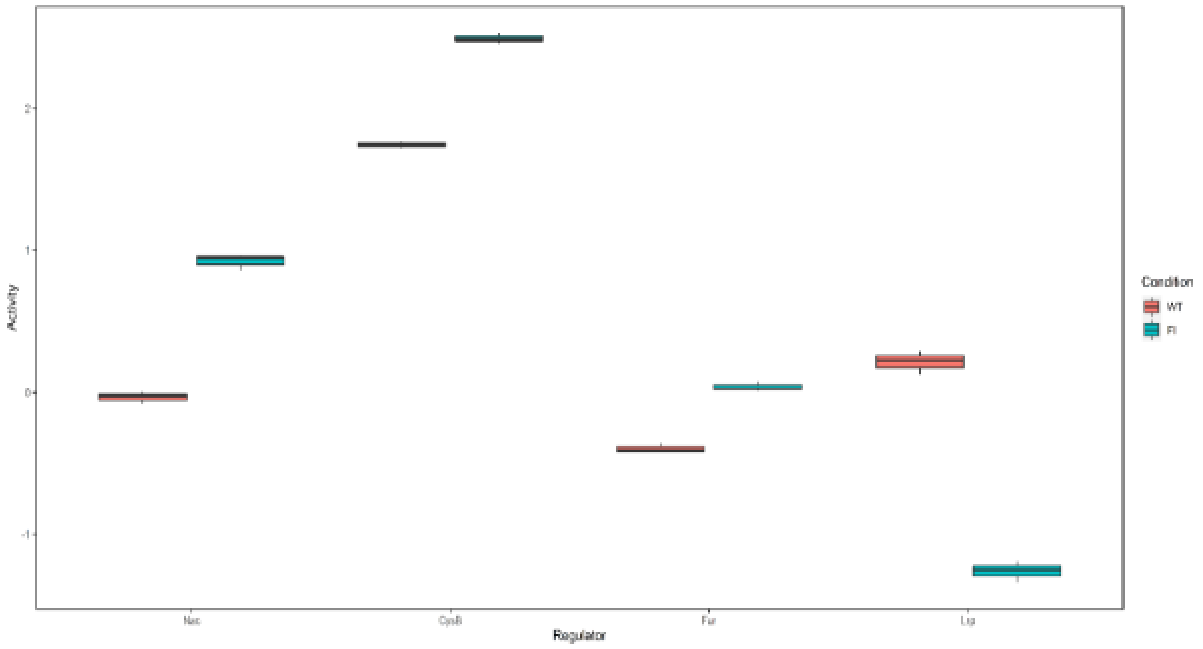


**F**

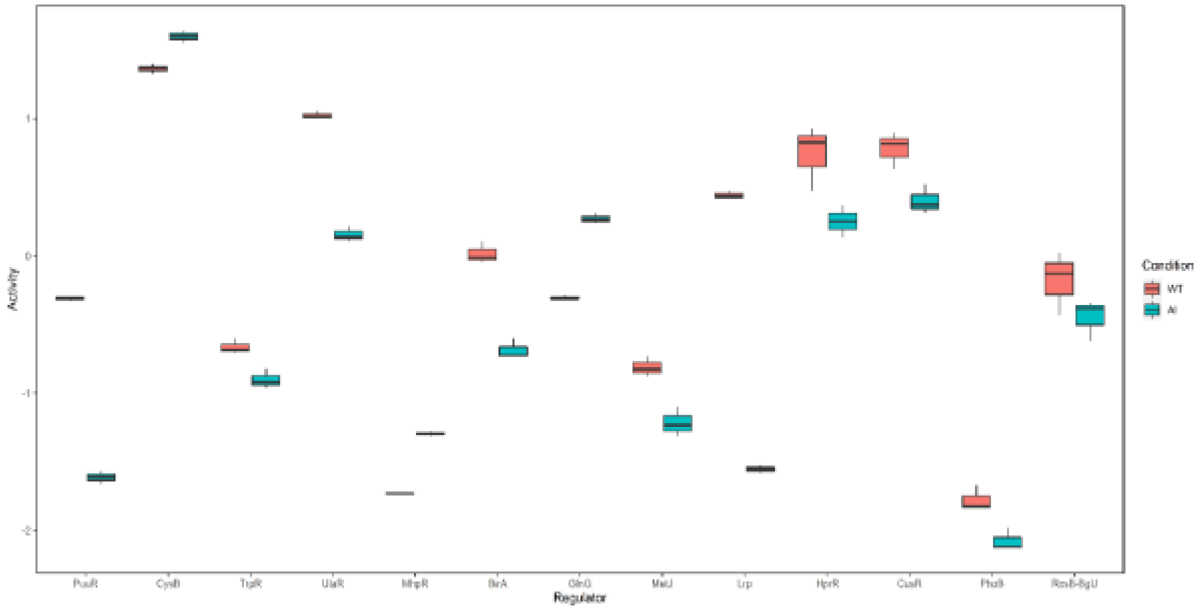


**Figure S7.** Boxplots representing regulatory activity (log-transformed) values of the iTFs in **A)** Δ*fnr*, **B)** Δ*arcA*, **C)** Δ*ihf*, **D)** FA, **E)** FI, and **F)** AI compared to WT. The activities were obtained from NCA (triplicate runs) as described in the methods section. Only the significantly enriched (FDR < 0.05) iTFs were used for the activity analysis.

**A**

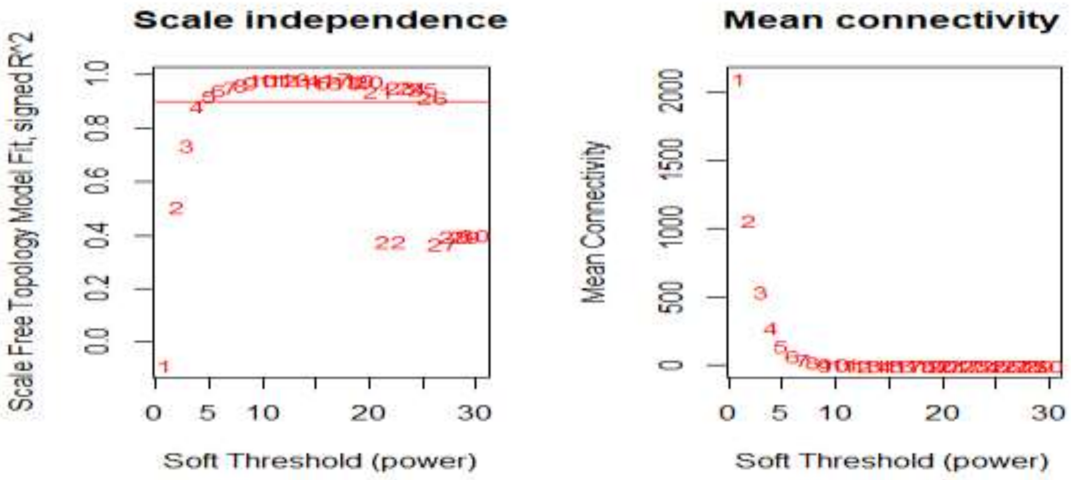


**B**

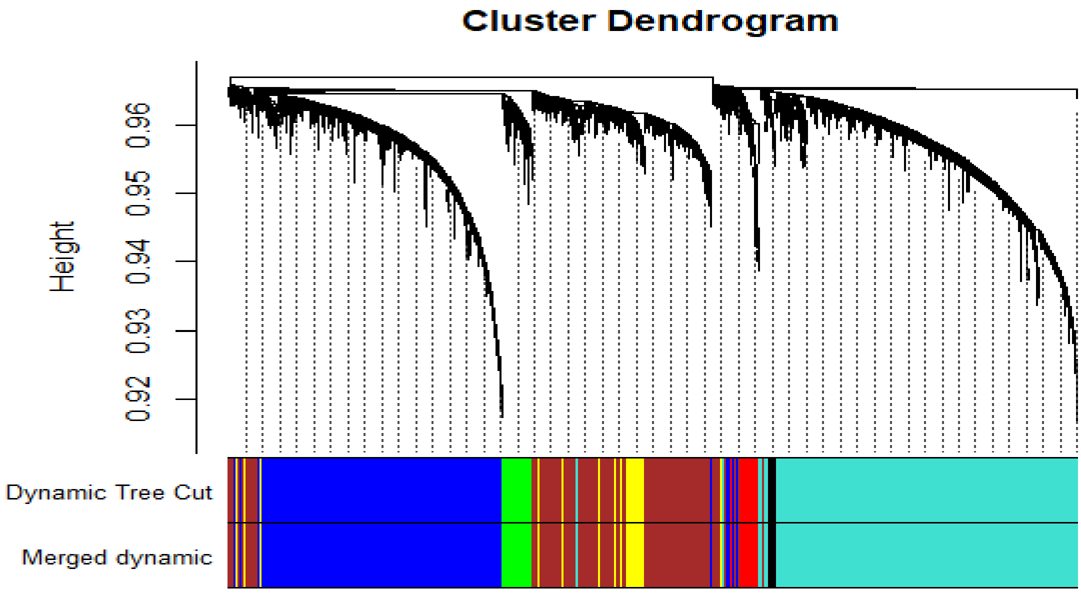


**Figure S8.** WGCNA analysis of *E. coli* K12 compendium gene expression data for construction of signed weighted co-expression networks. **A)** Soft thresholding to identify the power to achieve scale free topology index greater than 0.92. The soft threshold (power) was chosen to be 5 for our data with 4077 samples and 4189 genes. **B)** Clustering dendrogram considering the module eigengenes and dissimilarity value based on topological overlap. Each vertical line represents a single gene. The modules were assigned a colour and merged based on correlation value of ≥ 0.75.

A

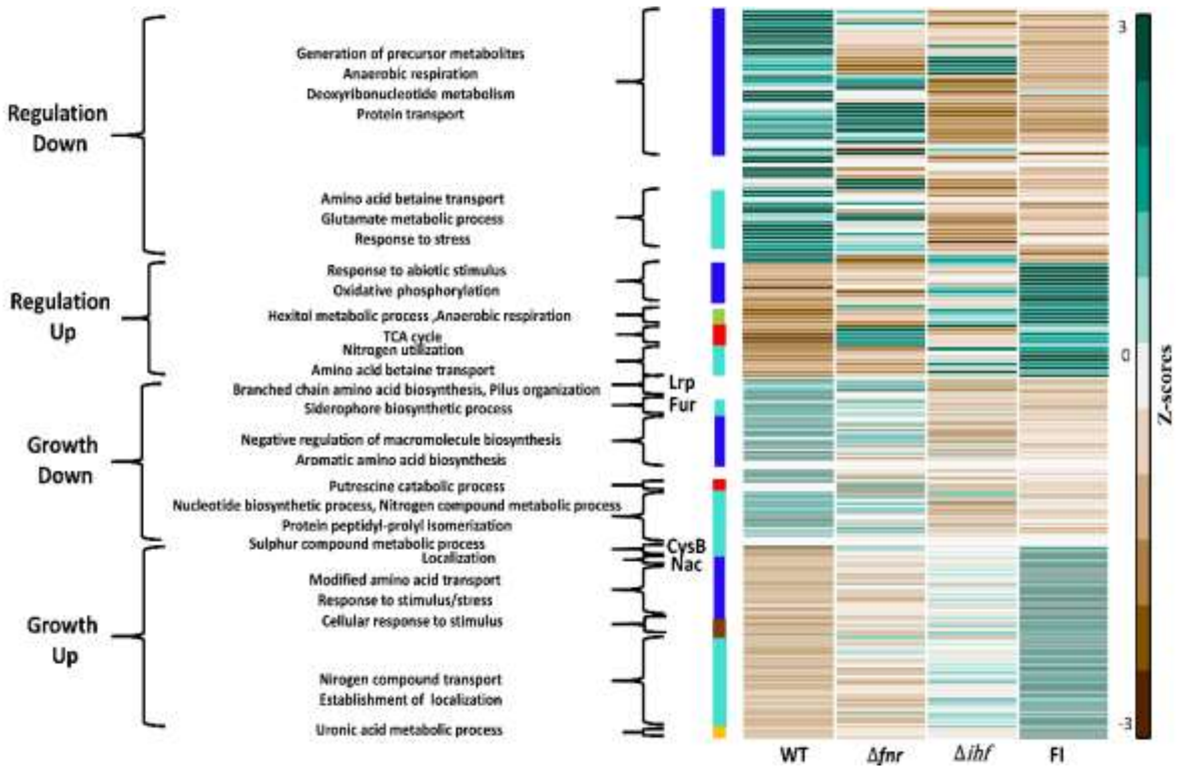
Regulation Down
Regulation Up
Growth Down
Growth Up
Generation of precursor metabolites
Anaerobic respiration
Deoxyribonucleotide metabolism
Protein transport
Amino acid betaine transport
Glutamate metabolic process
Response to stress
Response to abiotic stimulus
Oxidative phosphorylation
Hexitol metabolic process ,Anaerobic respiration
TCA cycle
Nitrogen utilization
Amino acid betaine transport
Branched chain amino acid biosynthesis, Pilus organization
Siderophore biosynthetic process
Negative regulation of macromolecule biosynthesis
Aromatic amino acid biosynthesis
Putrescine catabolic process
Nucleotide biosynthetic process, Nitrogen compound metabolic process
Protein peptidyl-prolyl isomerization
Sulphur compound metabolic process
Localization
Modified amino acid transport
Response to stimulus/stress
Cellular response to stimulus
Nirogen compound transport
Establishment of localization
Uronic acid metabolic process
Lrp
Fur
CysB
Nac
WT
Δfnr
Δihf
FI
3
0
-3
Z-scores

B

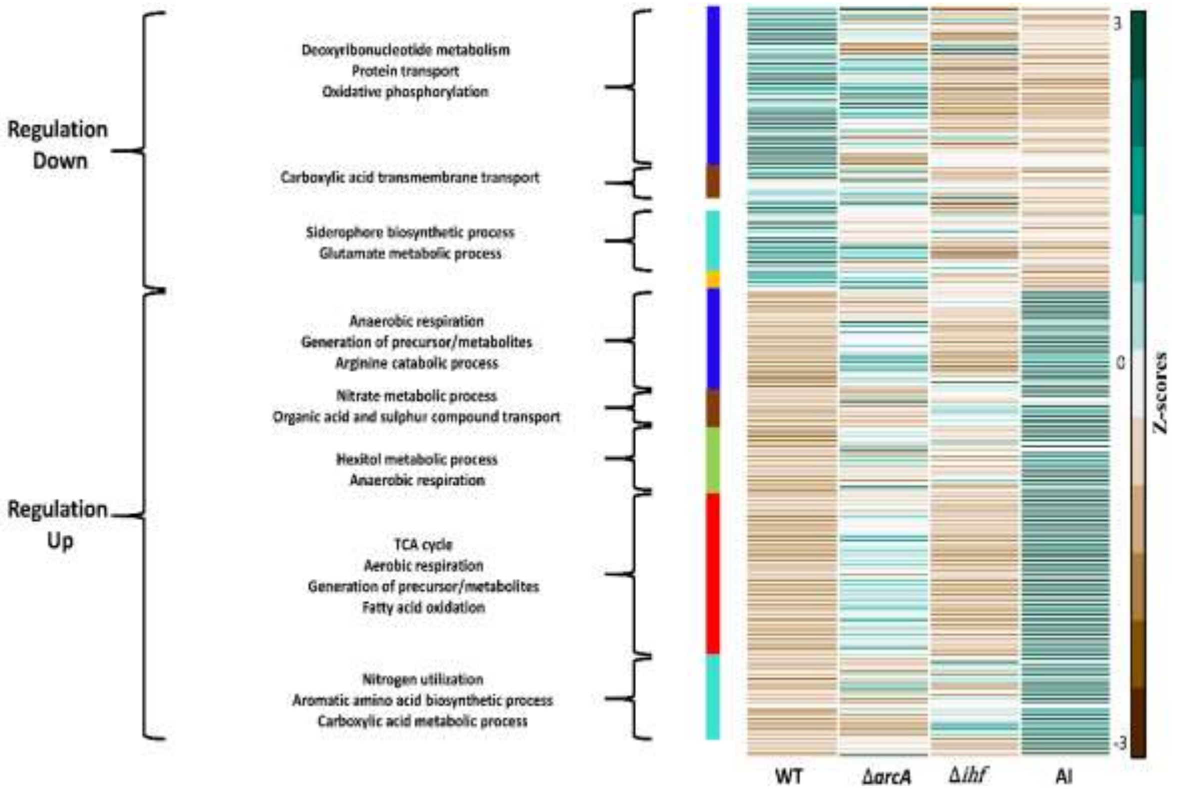
Regulation Down
Regulation Up
Deoxyribonucleotide metabolism
Protein transport
Oxidative phosphorylation
Carboxylic acid transmembrane transport
Siderophore biosynthetic process
Glutamate metabolic process
Anaerobic respiration
Generation of precursor/metabolites
Arginine catabolic process
Nitrate metabolic process
Organic acid and sulphur compound transport
Hexitol metabolic process
Anaerobic respiration
TCA cycle
Aerobic respiration
Generation of precursor/metabolites
Fatty acid oxidation
Nitrogen utilization
Aromatic amino acid biosynthetic process
Carboxylic acid metabolic process
WT
ΔarcA
Δihf
AI
3
0
-3
Z-scores

C

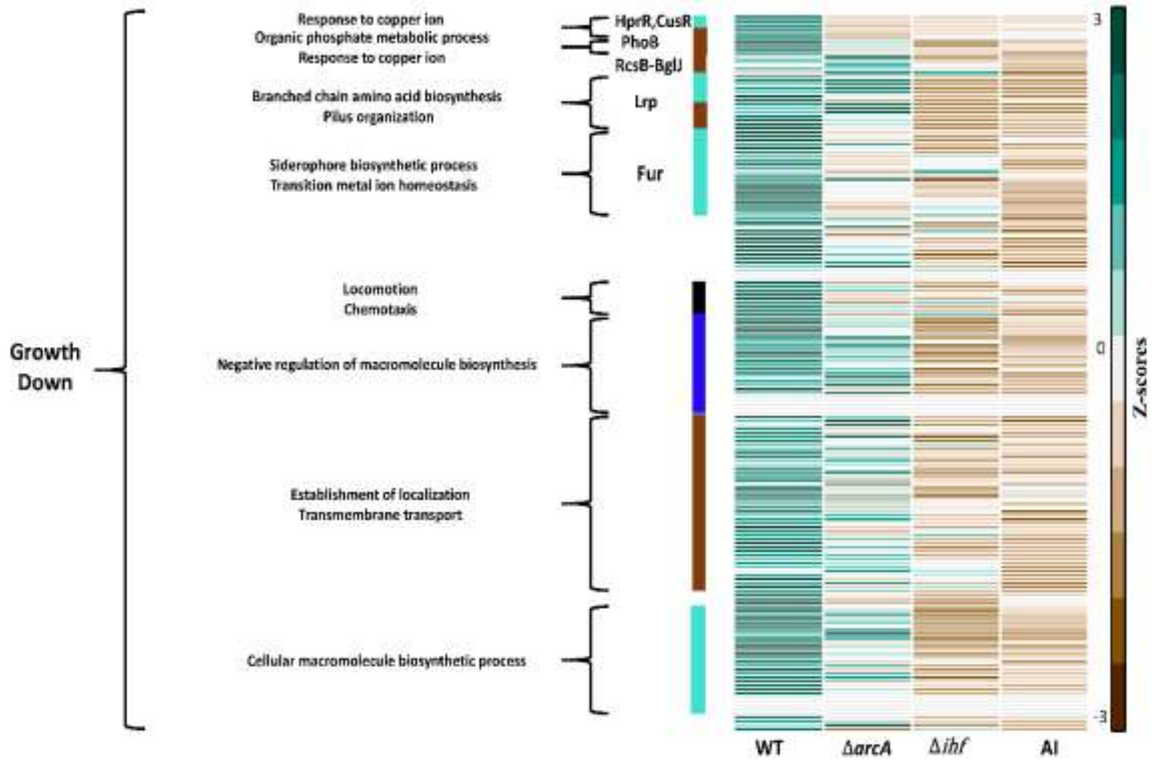
Response to copper ion
Organic phosphate metabolic process
Response to copper ion
Branched chain amino acid biosynthesis
Pilus organization
Siderophore biosynthetic process
Transition metal ion homeostasis
Locomotion
Chemotaxis
Growth Down
Negative regulation of macromolecule biosynthesis
Establishment of localization
Transmembrane transport
Cellular macromolecule biosynthetic process
HprR,CusR
PhoB
RcsB-BglJ
Lrp
Fur
3
0
-3
Z-scores
WT
ΔarcA
Δihf
AI

D

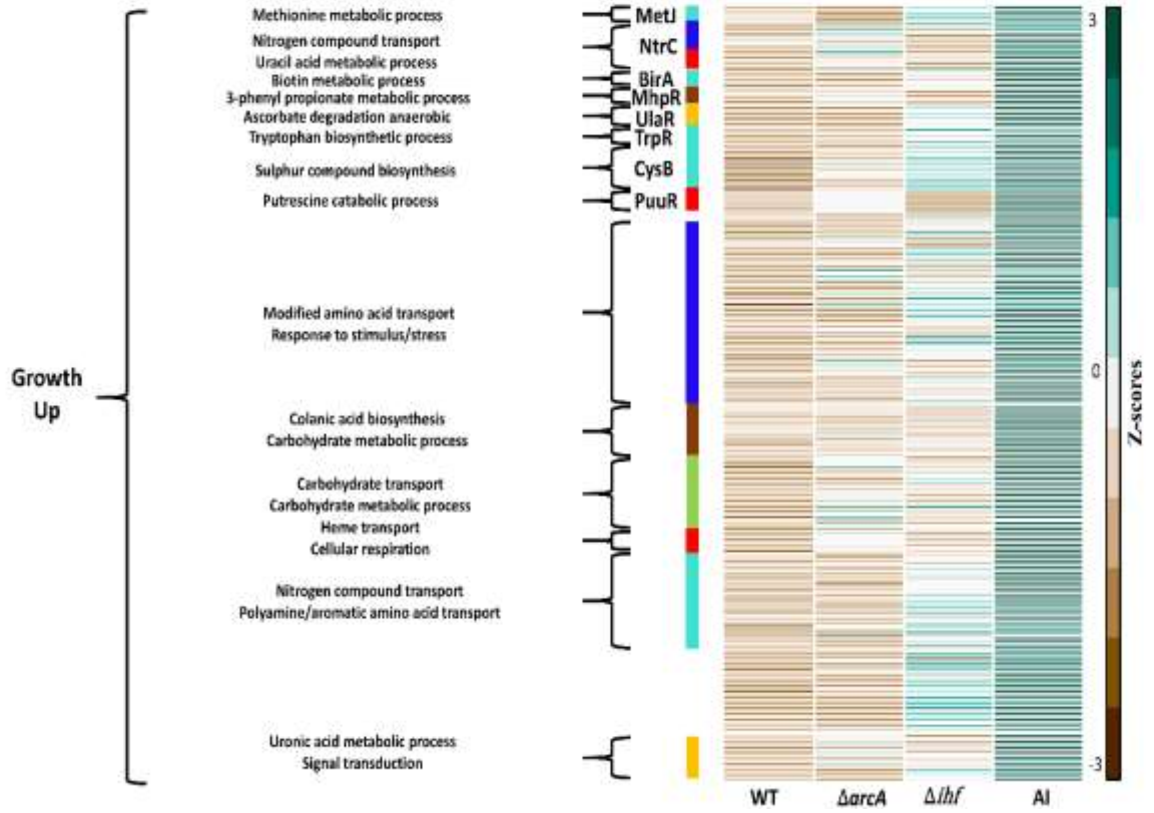
Methionine metabolic process
Nitrogen compound transport
Uracil acid metabolic process
Biotin metabolic process
3-phenyl propionate metabolic process
Ascorbate degradation anaerobic
Tryptophan biosynthetic process
Sulphur compound biosynthesis
Putrescine catabolic process
Modified amino acid transport
Response to stimulus/stress
Growth Up
Colanic acid biosynthesis
Carbohydrate metabolic process
Carbohydrate transport
Carbohydrate metabolic process
Heme transport
Cellular respiration
Nitrogen compound transport
Polyamine/aromatic amino acid transport
Uronic acid metabolic process
Signal transduction
MetJ
NtrC
BirA
MhpR
UlaR
TrpR
CysB
PuuR
3
0
-3
Z-scores
WT
ΔarcA
Δihf
AI

**Figure S9.** Heatmap depicting the regulation-specific direct and growth-rate mediated indirect targets for **A)** FI compared to WT and **B-D)** AI compared to WT. Both the upregulated and downregulated KPE DEGs in the FI and AI mutants compared to WT were used to compare across the corresponding single mutant strains. The tpm (transcript per million) values calculated for each of the double mutant and corresponding single mutant strains were z-score transformed. The colour strips next to the heatmap represent the different co-expression modules displayed here as different colors. The GO categories or TFs that have genes belonging to more than 1 module colour are shown as blank regions between the coloured strips.

### Text S1

The metabolite-TF interactions were considered plausible if the metabolite represents the product or substrate or part of any reactions within the metabolic subsystem of which the TF regulates the corresponding gene expressions. Perturbation in ArcA regulator activity across the strains were majorly involved in changes in gene expression of the putrescine catabolic process. Hence, we observed the association of metabolites such as glutamate, arginine and GABA due to their role in putrescine degradation. Such analogies can be extended to Crp and PuuR regulator activities. Modulation of Fur regulator activity across the strains involved changes in genes related to siderophore biosynthetic process or glycolysis (*gpmA*). The main compound for enterobactin biosynthesis is chorismate, which is derived from phosphoenolpyruvate. The increase in PEP modulates the Fur to repress the enterobactin biosynthesis so that PEP is made available for necessary biosynthetic processes. Similarly, PEP act as a precursor for aromatic amino acid biosynthesis and could potentially regulate TrpR activity to control tryptophan biosynthesis. The NtrC/GlnG regulator controls gene expression in response to nitrogen starvation. The metabolites αKG and glutamate play an important role in ammonia assimilation and hence could potentially modulate the activity of NtrC. Further, αKG and glutamate are crucial in branched chain amino acid synthesis, whose gene expression is regulated by Lrp. As aspartate is a key metabolite needed for synthesis of methionine, its association with MetJ seems reasonable. Glutamate is a structural analog to O-acetyl L-serine due to which we considered the occurrence of an association of CysB with glutamate.

| Mutants | Growth rate | Glucose uptake rate | Yield of Ethanol | Yield of Formate | Yield of Acetate | Yield of Lactate | Yield of Succinate | Yield of Pyruvate | Yield of Biomass |
|---|---|---|---|---|---|---|---|---|---|
| **WT** | 0.38 ± 0.007 | 16.63 ± 0.5 | 0.18 ± 0.01 | 0.35 ± 0.01 | 0.21 ± 0.01 | 0.02 ± 0 | 0.09 ± 0.01 | 0.02 ± 0 | 0.13 ± 0 |
| **IHF** | 0.29 ± 0.007 | 13.26 ± 0.23 | 0.19 ± 0 | 0.36 ± 0.01 | 0.23 ± 0 | 0.04 ± 0.017 | 0.07 ± 0.03 | 0.003 ± 0 | 0.12 ± 0 |
| **FNR** | 0.29 ± 0.008 | 12.12 ± 0.62 | 0.18 ± 0.01 | 0.39 ± 0.02 | 0.23 ± 0 | 0.04 ± 0 | 0.05 ± 0 | 0.002 ± 0 | 0.13 ± 0.01 |
| **ArcA** | 0.32 ± 0.015 | 13.99 ± 0.44 | 0.18 ± 0.005 | 0.33 ± 0.02 | 0.21 ± 0.01 | 0.01 ± 0 | 0.12 ± 0.01 | 0.002 ± 0 | 0.13 ± 0 |
| **FA** | 0.23 ± 0.002 | 11.27 ± 0.43 | 0.17 ± 0.007 | 0.36 ± 0.005 | 0.23 ± 0.009 | 0.04 ± 0.01 | 0.09 ± 0.004 | 0.002 ± 0 | 0.11 ± 0.005 |
| **FI** | 0.273 ± 0.011 | 13 ± 0.5 | 0.18 ± 0.004 | 0.37 ± 0.003 | 0.24 ± 0.002 | 0.05 ± 0.011 | 0.07 ± 0.008 | 0.001 ± 0 | 0.12 ± 0.002 |
| **AI** | 0.26 ± 0.005 | 14.15 ± 0.32 | 0.2 ± 0.008 | 0.32 ± 0.004 | 0.21 ± 0.005 | 0.01 ± 0 | 0.17 ± 0.002 | 0.004 ± 0 | 0.1 ± 0 |

**Table S1.** Physiological characterization of the strains in anaerobic fermentation of glucose. The measurements of growth rate ($h^{-1}$), glucose uptake rate (mmol/gDCW/h) and yields (g/g glucose) of mixed-acid fermentation products were obtained from three biological replicates ($n = 3$). Yields were calculated by normalizing secretion rates with its glucose uptake rate. The errors indicate standard deviations within the replicates. The measurements for the single mutants were obtained from our previous study (Iyer et al., 2021).

| **Strain Name** | **Genotype** | **Source** |
|---|---|---|
| *E. coli* K-12 MG1655 WT | F-, *λ-*, *ilvG- rfb-50 rph-1* | Keio collection CGSC #6300 |
| *E. coli* K-12 MG1655 Δ*arcA* | F-, *λ-*, *ilvG- rfb-50 rph-1, ΔarcA::kan* | Iyer et al., 2021 |
| *E. coli* K-12 MG1655 Δ*fnr* | F-, *λ-*, *ilvG- rfb-50 rph-1,* Δ*fnr::kan* | Iyer et al., 2021 |
| *E. coli* K-12 MG1655 Δ*ihf* | F-, *λ-*, *ilvG- rfb-50 rph-1,* Δ*ihfA::FRT* Δ*ihfB::kan* | Iyer et al., 2021 |
| *E. coli* K-12 MG1655 FA | F-, *λ-*, *ilvG- rfb-50 rph-1,* Δ*fnr::FRT ΔarcA::kan* | This study |
| *E. coli* K-12 MG1655 FI | F-, *λ-*, *ilvG- rfb-50 rph-1,* Δ*fnr::cm* Δ*ihfA::FRT* Δ*ihfB::kan* | This study |
| *E. coli* K-12 MG1655 AI | F-, *λ-*, *ilvG- rfb-50 rph-1,* Δ*arcA::cm* Δ*ihfA::FRT* Δ*ihfB::kan* | This study |

**Table S2.** Strains used in this study.
